# Mating and ecdysone signaling modify growth, metabolism, and digestive efficiency in the female *Drosophila* gut

**DOI:** 10.1101/2024.11.19.624434

**Authors:** Tahmineh Kandelouei, Madeline E. Houghton, Mitchell R. Lewis, Caroline C. Keller, Marco Marchetti, Xiaoyu Kang, Bruce A. Edgar

## Abstract

Adaptive changes in organ size and physiology occur in most adult animals, but how these changes are regulated is not well understood. Previous research found that mating in *Drosophila* females drives not only increases in gut size and stem cell proliferation but also alters feeding behavior, intestinal gene expression, and whole-body lipid storage, suggesting altered gut metabolism. Here, we show that mating dramatically alters female gut metabolism and digestive function. In addition to promoting a preference for a high-protein diet, mating also altered levels of TCA cycle intermediates and fatty acids in the gut, increased total gut lipids and protein, reduced relative carbohydrate levels, and enhanced the efficiency of protein digestion relative to carbohydrate digestion. The expression of genes that mediate each of these metabolic processes was similarly altered. In addition, we noted the mating-dependent downregulation of oxidative stress response and autophagy genes. Mating-dependent increases in ecdysone signaling played an important role in re-programming many, but not all, of these changes in the female gut. This study contributes to our understanding of how steroid signaling alters gut physiology to adapt to the demands of reproduction.

## INTRODUCTION

Adaptive changes in organ size and physiology are crucial for the survival and optimal health of adult animals. Such adaptive changes occur in response to various internal and external changes, including alterations in an organism’s physiological state, nutritional state, or environment. In mammals, adaptive growth of the intestine during pregnancy and lactation can be more than double to size of the organ, and this is crucial to meet the increased nutritional requirements^1-4^ of both the mother and her offspring. This requires a complex interplay of hormonal, growth factor, and nutritional signals that regulate structural and functional changes in the gastrointestinal tract. Although progress has been made in understanding these mechanisms, many aspects remain incompletely understood ^5-7^. Ongoing research aims to better understand the precise molecular mechanisms and regulatory pathways involved in adaptive remodeling of the intestine.

In females of *Drosophila melanogaster*, mating causes significant changes in behavior, including increased egg-laying frequency, decreased sexual receptivity, increased food consumption, decreased immunity, and shortened lifespan ^8-12^. These changes are associated with a shift in food preference towards foods with increased protein content ^13, 14^. The reproductive performance of female flies, especially egg production, is directly related to their nutritional status, with higher protein intake required for higher fecundity^15, 16^. Mating triggers a ∼60% enlargement of the female gut through the actions of 20-hydroxy-ecdysone (20HE) ^14, 17-19^, Sex Peptide (SP) ^14^, and possibly juvenile hormone (JH)^19^, which stimulate the proliferation of intestinal stem cells ^17, 18^. Like vertebrate estrogen and testosterone, ecdysone is essential for developmental transitions and sexual maturation in insects ^20^, and it also influences physiological processes, including metabolism and longevity in adults ^12, 21, 22^. However, mating-induced enlargement of the adult fly gut, mediated by 20HE, predisposes females to intestinal dysplasia and tumor formation ^17^. Transcriptomic and metabolomic analyses of whole flies have revealed a post-mating metabolic shift favoring protein utilization and lipid production, which was proposed to not only facilitate greater fecundity but to make females more susceptible to bacterial toxicity and to shorten their lifespan ^11^. Female flies require the hormone Sex Peptide (SP), found in male seminal fluid, to trigger gut enlargement and modulate the expression of digestive enzymes and metabolic genes after mating ^14^. Maternal food increase in mated flies through the enlargement of the crop in the anterior gut (an expandable structure in the insect intestine) is adaptive and regulated by Myosuppressin neurons. These neurons are targets of ecdysone, and have been shown to promote food intake^10^. All these findings from previous research inform the central question of this project, namely: what is the nature of mating-induced metabolic changes in the female gut, and how does ecdysone signaling regulate them? To answer this question, we investigated how mating affects gut physiology and growth, and in particular whether mating alters gut metabolism. We sought to identify the metabolic changes associated with intestinal growth, learn whether mating alters digestive function, and determine whether ecdysone regulates these processes.

## RESULTS

### Mating-induced gut growth alters gut cell- and chemical composition

We first sought to reproduce previous reports that mating promotes adaptive gut growth in females. In our experiments, mated female flies were cultured with males, whereas virgins were isolated apart from males from eclosion. We allowed mating to begin 4 days post-eclosion and allowed the flies to mate for two days at 25°C. After this 2-day mating period (6 days post eclosion), we dissected out the midguts and quantified the numbers of progenitor and total cells and gut areas by immunofluorescent staining and flow cytometry. Consistent with previous reports ^14, 17-19^, mated females had larger intestinal areas in the R2 and R4 regions (Figure 1A, 1B) and approximately 73% more *esg+* progenitor cells (ISC and EB; Figure 1C). The total number of cells assayed by fluorescence-activated cell sorting (FACS) of isolated nuclei was also increased by approximately 57% (Figure 1G). The distributions of ploidy values of the nuclei were not significantly changed, however, with nuclei from mated and virgin intestines showing similar distributions of nuclei ranging from 2C to 64C in DNA content (Figure 1H). Since DNA ploidy values typically parallel cell sizes quite closely, this indicates that mating-dependent gut growth is driven primarily by increases in cell numbers rather than by increased cell sizes. This is consistent with previous studies that reported increased intestinal stem cell (ISC) divisions after mating and minimal enlargement of mature enterocytes ^17, 18^.

**Figure 1.**
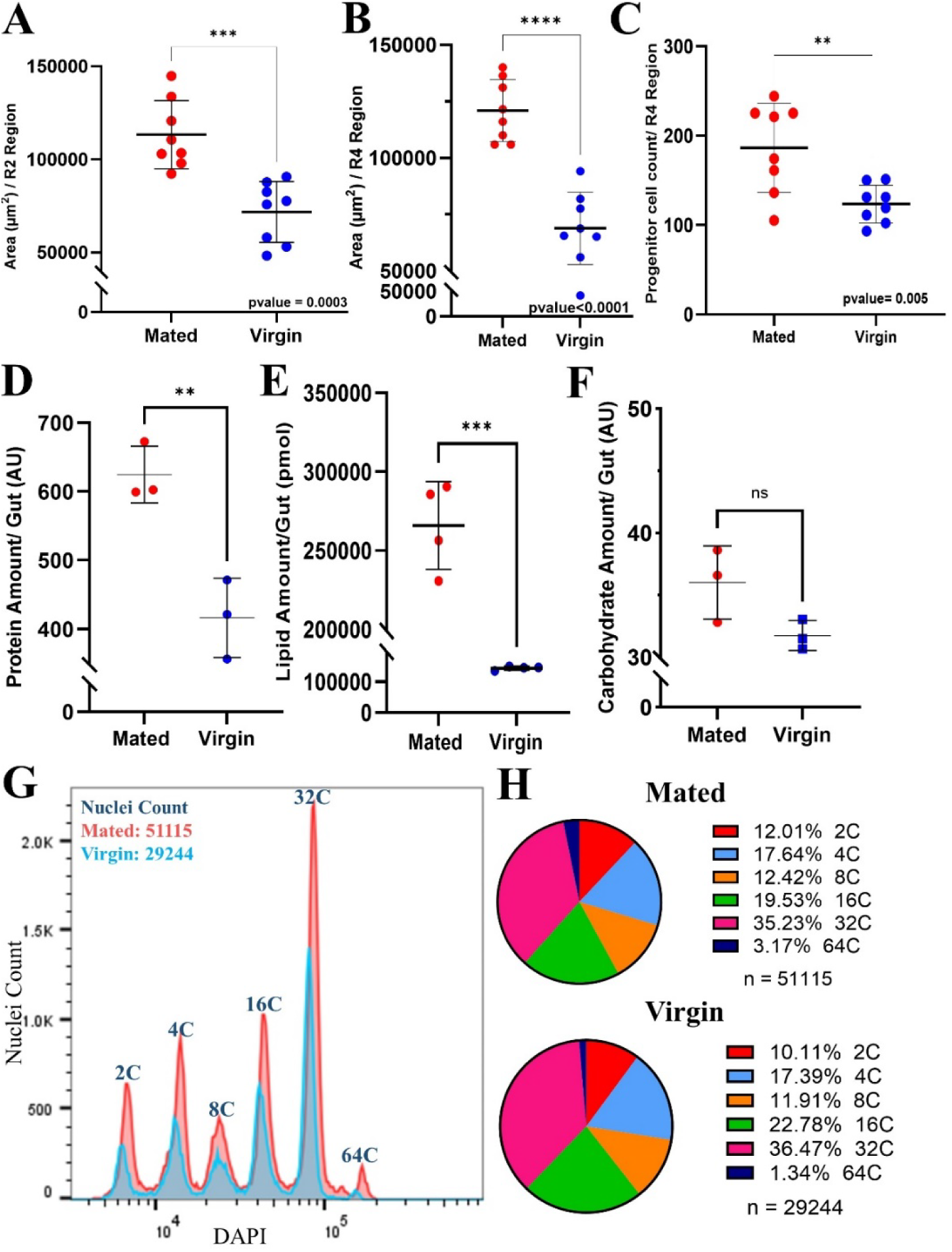
Mating induces region-specific adaptive gut growth and alters overall gut composition. **A.** R2 region area measurement of mated and virgin flies (*esg-Gal4^ts^*). **B.** R4 region area measurement of mated and virgin flies (*esg-Gal4^ts^*). **C.** Progenitor cell count in the R4 region of mated and virgin flies (*esg-Gal4^ts^*). **D.** Total protein content/gut defined by BCA assay comparing mated and virgin flies defined by colorimetric assay in mated (3 replicates/15 guts) and virgin flies (3 replicates/15 guts) (*Tub-Gal4^ts^*). **E.** Total lipid content (pmol) defined by LC-MS of mated flies (3 replicates/40 guts) and virgin flies (3 replicates/ 40 guts) (*esg-Gal4^ts^*). **F.** Total carbohydrate content/gut defined by colorimetric assay in mated (3 replicates/15 guts) and virgin flies (3 replicates/15 guts) (*Tub-*Gal4^ts^). **G.** Histogram depicting FACS-sorted nuclei from mated and virgin fly midguts, using 25 midguts/sample (*esg-Gal4^ts^*); mating was for 5 days. **H.** Ploidy distributions of FACS-sorted midgut nuclei from the samples in G.

Next, we sought to learn how mating affects the composition of macromolecules, including proteins, lipids and carbohydrates in the gut. Total gut protein content, as determined by a colorimetric assay, showed very significant (∼60%) increases after mating (Figure 1D), paralleling the increase in total cell numbers. Likewise, lipidomics analyses by mass spectrometry showed large (∼100%) increases in total gut lipid amounts in mated flies (Figure 1E). However, quantification of total gut carbohydrate contents using a colorimetric assay showed that mated flies had relatively small increases in carbohydrate contents that were not statistically significant (Figure 1F). Comparisons of the relative changes in gut components after mating showed that the increase in lipid content (∼100%) was relatively much greater than the increase in protein (∼60%) or carbohydrate (∼12%; Figure 1D-F). These changes in the relative amounts of macromolecules likely reflect major alterations in gut cell physiology and function.

### Mating alters gut TCA cycle metabolism

Since mating changes overall gut composition, we investigated how it alters the gut metabolic profile. Steady-state metabolite profiling on isolated midguts by mass-spectrometry showed significant mating-dependent increases in the gut content of specific fatty acids, carbohydrates, amino acids, and other small metabolites. In particular, metabolite set enrichment analysis showed significant changes in metabolites of the tricarboxylic acid cycle (TCA), amino acid metabolism, and intermediates in fatty acid and carbohydrate metabolism (Figure 2A) (Extended Figure 5A). Increased TCA cycle intermediates included fumaric acid (log2FC: 0.55, p-value< 0.05), malic acid (log2FC: 0.85, p-value< 0.05) and myo-inositol (log2FC: 0.64, p-value< 0.05; Extended Figure 4A, Data S1). TCA cycle metabolites have long been regarded as by-products of cellular metabolism that are essential for both energy (NADH, ATP) production and the biosynthesis of macromolecules, including nucleotides, lipids, and proteins. While these functions are crucial for the maintenance of cellular homeostasis, it is increasingly recognized that TCA cycle metabolites also play important roles in regulating chromatin modifications, DNA methylation, and post-translational modifications of proteins, thereby altering gene functions ^23, 24^.

**Figure 2.**
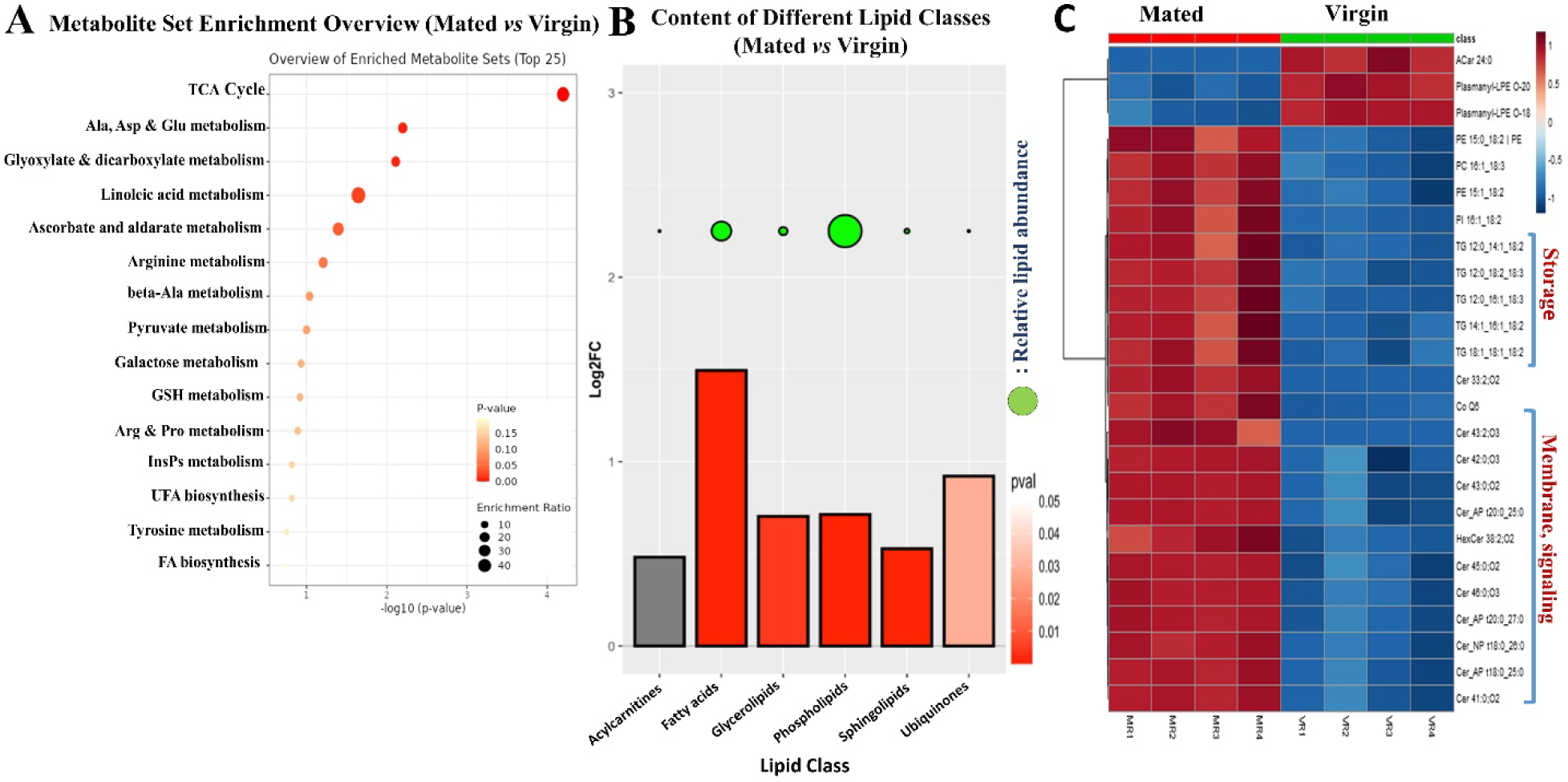
Mating increases TCA cycle metabolites, TGs, ceramides, fatty acids, carbohydrates, and amino acids content in the gut. **A.** GC-MS metabolic set enrichment overview in mated flies (*Tub-Gal4^ts^*) (4 replicates/60 guts) compared to virgin controls (*Tub-Gal4^ts^*) (4 replicates/ 60 guts). **B.** LC-MS analysis-derived bar graphs show the enrichment of different classes of lipids in mated flies gut (*esg-Gal4^ts^*) (4 replicates/ 40 guts), compared to virgin (*esg-Gal4^ts^*) (4 replicates/40 guts), the color of each bar shows the significant of the change, red shows statistical significant change while gray color shows insignificant change, the y-axis shows the log-2 fold changes, and x-axis shows the values, green disc size shows the abundance of its correspondent class of lipid in the graph. **C.** LC-MS analysis shows the top forty lipid species (comparing mated and virgin flies gut) with log-2 fold changes>1 and significant p-values (< 0.05) depicted in the heat map based on z-score.

### Mating increases intestinal triglycerides and ceramides

As noted above, mating also increased the total gut lipid content and the relative fraction of lipids in the gut (Figure 1E). A detailed analysis by liquid chromatography-mass spectrometry (LC-MS) showed that mating-induced increases in lipids varied by class (Figure 2B). Triglycerides and ceramides showed the greatest, most significant increases among all detected lipid species (Figure 2C). Gas chromatography-mass spectrometry (GC-MS) data showed that fatty acids, especially linoleic acid (log2FC=0.72, p< 0.05) and lauric acid (log2FC=1.0, p< 0.05) were also increased in the guts of mated flies, suggesting changes in fatty acid synthesis and/or utilization (Extended Figure 4A). Linoleic acid is a major component of phospholipids in cell membranes and is essential for successful egg production and viability ^25^. It can also be metabolized for energy, which is critical during times of high metabolic demand or rapid growth, such as when it is required for efficient egg production. Similarly, lauric acid can be used either for the synthesis of more complex lipids, which were dramatically increased in the gut after mating, or metabolized for energy. As the longest of the medium-chain fatty acids, lauric acid can be transported into mitochondria either directly or through the carnitine shuttle system as a fast source for β-oxidation, Acetyl-CoA, and ATP. This is particularly important for physiological states with high-energy requirements, such as reproduction. Indeed, acylcarnitine 12:0 associated with lauric acid, the form which is shuttled into mitochondria, was more than twice as abundant in mated flies than virgins (33.7 *vs* 14.7 pMol) (See Data S1). Overall, these results emphasize physiological changes induced by mating in female flies. The guts in mated animals are not only larger but are qualitatively different in as much as the proportions of molecular components are changed. The many metabolite-specific changes we observed, including in TCA cycle intermediates, amino acids, fatty acids, triglycerides, and ceramides, indicate major alterations in the regulation and function of metabolic pathways during mating-dependent gut remodeling.

### Mating alters mRNA expression profiles in gut progenitor and differentiated cells

Previous studies reported major changes in intestinal gene expression in response to sex determination ^26^, as well as mating, in females ^14^. To learn how mating drives adaptive gut growth and remodels gut physiology and metabolism, we performed RNAseq analyses on both FACS-sorted *esg+* progenitor cells and whole guts from mated and virgin females. RNAseq profiles taken at 48 hours after mating showed significant changes in the levels of mRNAs for genes involved in many metabolic processes, including but not limited to: upregulation of *Jonah*-type serine endopeptidases used in protein digestion (*Jon99Fii, Jon99Cii, Jon44E, Jon65Ai, Jon25Bi, Jon25Bii, Jon99Ciii, Jon65Aiv, Jon65Aiii*) (Extended Figure 1A); upregulation of genes encoding amino acid transporters (*dmGlut, slif, NAAT1*); and relative downregulation of maltases and other genes involved in carbohydrate catabolism (*Mal-A4*, *Mal-A7, Mal-A8, Mal-6, Mal-1, Mal-2*). RNAseq analyses of FACS-sorted progenitor cells showed similarities to these trends in the whole gut and, in addition, revealed a progenitor cell-specific mating-dependent downregulation of *Gst* family genes (*GstD2, GstD5, GstD6, GstD9, GstE1, GstE3, GstE6, GstE7, GstE8, GstE9, GstO3, GstT4*), which are involved in combating oxidative stress (Extended Figure 1A); for the full list of all genes, see also Data S2. We used Gene Ontology (GO) enrichment analysis to group genes based on their functions in defined GO terms (Whole gut RNAseq) (Figure 3A, B). GO enrichment analysis of all the genes that were changed most significantly showed that mating causes a general upregulation of genes involved in proteolysis (Figure 3A) and downregulation of genes involved in carbohydrate metabolic processes (Figure 3B). Gene set enrichment analysis (GSEA) also confirmed that mating upregulates the expression of proteolysis genes (Extended Figure 1B) while it downregulates carbohydrate metabolic process genes (Whole gut RNAseq) (Extended Figure 1C). GO enrichment analysis of whole gut RNAseq also revealed the mating-dependent downregulation of the majority of autophagy-related genes, including, for instance, *Atg1*, *Atg2*, *Atg3*, *Atg4b*, *Atg9*, *Atg18a*/*b, Atg101* and *Rab18* (GO:0000045 and GO:0005776) (Figure 3F). Interestingly, this trend was not observed in *esg+* progenitor cells, which, in fact, showed a reverse trend (Extended Figure 3A). See also Data S2 and Data S3.

**Figure 3.**
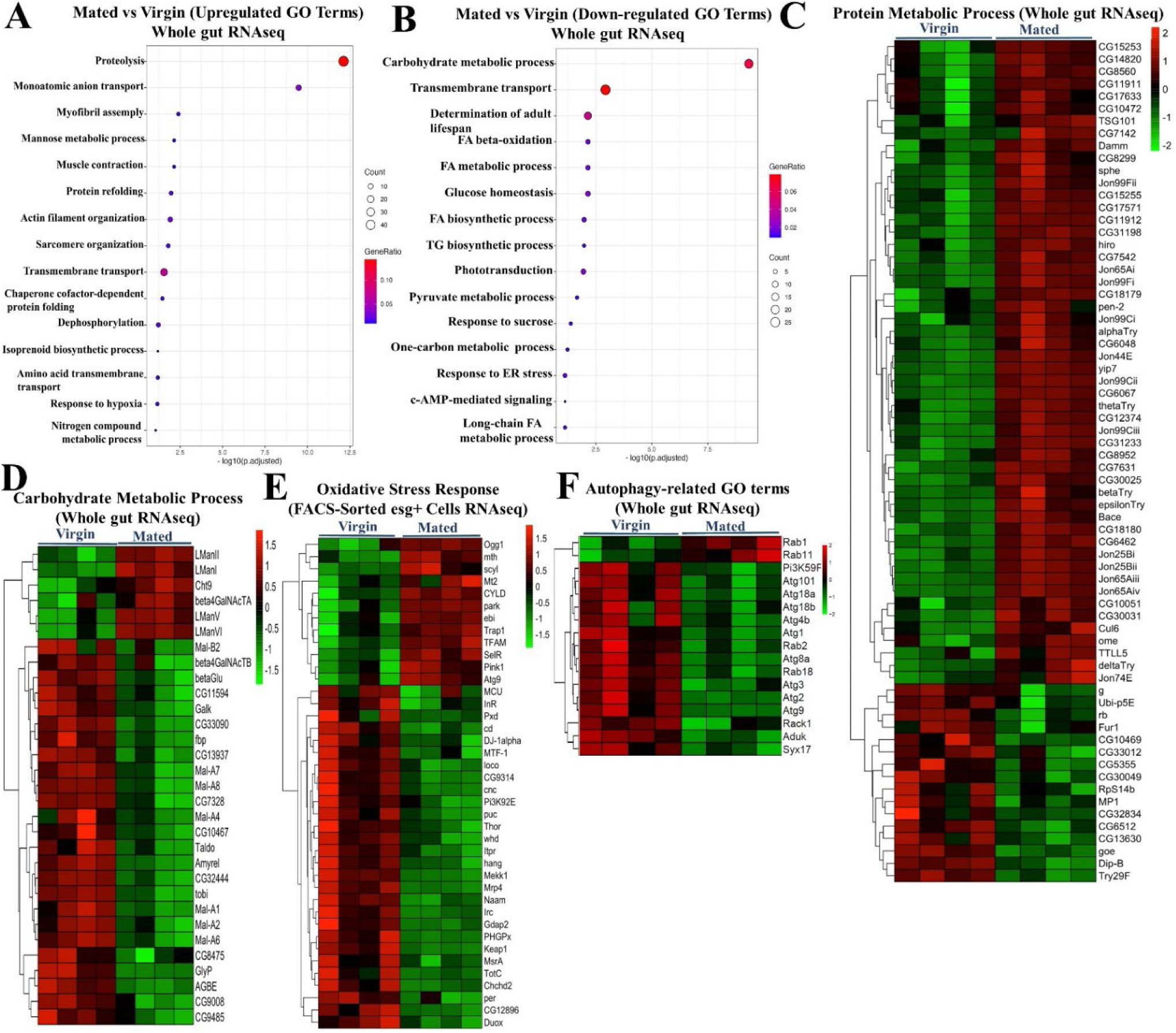
Mating upregulates protein metabolism gene expression and downregulates carbohydrate metabolism and oxidative stress response and autophagy genes expression. **A.** GO enrichment analysis showing upregulated pathways in guts of mated flies (*esg-Gal4^ts^*) compared to virgin ones (*esg-Gal4^ts^*) (each/4 replicates/15 guts) (Whole gut RNAseq). **B.** GO enrichment analysis showing downregulated pathways in guts of mated flies (*esg-Gal4^ts^*) compared to virgin controls (*esg-Gal4^ts^*) (each/4 replicates/15 guts) (Whole gut RNAseq). **C-F.** Targeted heat maps for the significantly differentially expressed genes classified under the GO terms “protein metabolic process” (whole gut RNAseq; C), “carbohydrate metabolic process” (whole gut RNAseq; D), “oxidative stress response metabolic process” (FACS-sorted esg+ cells RNAseq; E) and “autophagy-related GO terms” (whole gut RNAseq; F).

In addition, we noted another set of genes with significant differential expression, namely genes involved in stress responses (FACS-sorted esg+ cells RNAseq) (Extended Figure 1D), most notably oxidative stress response genes (Extended Figure 1E). Most of these genes were downregulated in the gut by mating. Notably, a previous study ^14^ uncovered these same classes of genes as altered in the gut by mating and showed additionally that their induction or repression was dependent upon male-derived Sex Peptide.

To gain more insight into the regulation of the metabolic pathways changed by mating, we considered our metabolite and RNAseq profiling data sets together. Lipidomics analysis clearly showed increases in gut triglycerides (TG), storage of form of lipid, and RNAseq showed significant upregulation of genes such as the triacylglycerol lipase *brummer (bmm)* (Extended Figure 1A), which is used in mobilizing stored (and perhaps dietary) fats ^27^. GC-MS showed increases in fatty acids, which are the precursors for TG production. Our GC-MS metabolomics data also showed significant changes in amino acids, including arginine, tryptophan, and valine, which all increased after mating. Looking into gene expression data sets showed the upregulation of the *Jonah* genes, which encode gut-specific serine-type endopeptidases that play a central role in the digestion of dietary proteins by breaking them down into smaller peptides and eventually amino acids for absorption. They show homology with mammalian serine proteases such as trypsin and chymotrypsin. Together, these results suggest that mating in female *Drosophila* melanogaster leads to metabolic alterations in the gut that favor the utilization of proteins and fats over the processing of carbohydrates, probably to meet the increased energy demands and biosynthesis processes associated with reproduction (egg production).

### Mating-dependent changes in gut lipids and carbohydrates are ecdysone signaling-independent

Next, we sought to understand whether any of the mating-induced changes in metabolite and transcriptome profiles were due to ecdysone signaling, which we and others previously identified as instructive for promoting mating-dependent gut growth and female fecundity ^17, 18^. To investigate how the ecdysone signaling pathway affects mating-induced changes, we compared mated and virgin flies, each expressing or not expressing *EcR-RNAi*. We aged the collected virgin flies for 5 days at 29°C to express *EcR-RNAi* under the control of ubiquitous (*Tubulin-Gal4; Tub-Gal4)* or EC-specific (*Myo1A-Gal4)* drivers. After 5 days of RNAi expression, we set the mating for 48 hours (25°C) by adding males. We then dissected the guts after 48 hours of mating for sample preparation. While previous studies^17, 18^ had focused on EcR-dependent functions in intestinal progenitor cells (ISC, EB) and used the progenitor cell-specific *esg-Gal4* driver, we employed conditional (temperature-sensitive) versions of the ubiquitously expressed *Tubulin-Gal4 (Tub-Gal4^ts^)* and EC-specific *Myosin1A-Gal4* (*Myo1A^ts^*) drivers to assess potential EcR dependent functions in other cell types.

As shown above (Figure 1E), mating increased the total lipid content of the *Drosophila* female intestine. Phospholipids, triglycerides, and fatty acids were the major classes of lipids that accounted for the bulk of the mating-dependent lipid increase (Figure 2B), and ceramides and triglycerides showed the most significant increases among LC-MS detected lipid species (Figure 2C). Yet, in tests using *EcR-RNAi*, LC-MS lipidomics analysis revealed that the total increase in lipid content in response to mating was not significantly changed when ecdysone signaling was inhibited (Figure 4A). As in controls, the bulk of the lipid increase could be attributed to increased triglycerides, a storage form of lipid in lipid droplets in ECs. As noted above, our assays showed that mated flies did not have significantly more carbohydrates in their guts than virgins (Figure 1F). Likewise, further analysis showed that *EcR-RNAi* had no detectable effect on gut carbohydrate content (Extended Figure 4B).

**Figure 4.**
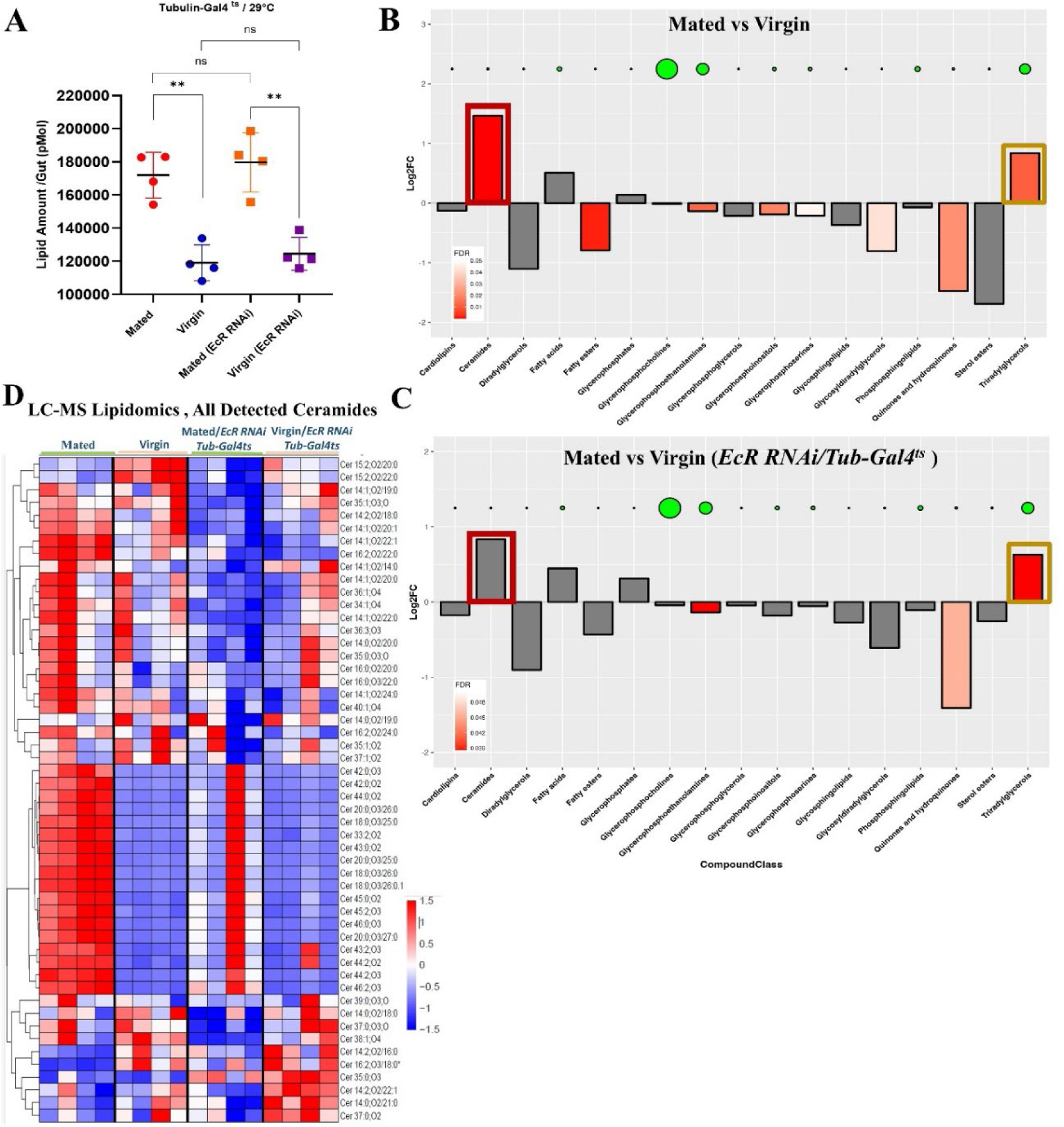
Mating-induced increase in ceramides is EcR-dependent. **A.** Total lipid content derived from LC-MS comparing mated and virgin flies +/- *EcR-RNAi* using *Tub-Gal4*^ts^, each/ 4 replicates/ 40 guts. **B.** LC-MS derived bar graphs show enrichment of different classes of lipids in mated flies compared to virgin flies; green disk size shows the abundance of the correspondent lipid class in the graph. **C.** LC-MS derived bar graphs show enrichment of different classes of lipids in mated flies with *EcR-RNAi* expression (*Tub-Gal4^ts^*) compared to genotype-matched virgin flies; green disk size shows the abundance of the correspondent lipid class in the graph. **D.** Targeted heatmap from LC-MS data analysis shows all detected ceramide species comparing mated and virgin flies gut +/- *EcR-RNAi* using *Tub-Gal4^ts^* depicted in the heat maps based on z-score (each/4 replicates/ 40 guts).

### Mating-induced increases in ceramides in the gut are ecdysone signaling-dependent

In contrast to bulk lipids (Figure 4A) and the major lipid classes (Figure 4B, C), we did note that the mating-dependent enrichment of most detected ceramide variants was significantly ablated by *EcR-RNAi* (Figure D, see also Data S1). Ceramides are known to induce the proliferation of ISCs by promoting the utilization of fatty acids ^28^. As mating-induced ISC proliferation also depends on ecdysone signaling ^17, 18^, this correlation suggests that *EcR-RNAi* might suppress ISC proliferation by interfering with ceramide metabolism.

### Mating-induced increases in total gut protein are ecdysone signaling-dependent

Whereas mating increased the total protein content of female fly gut (Figure 1D), the guts of mated flies expressing *EcR-RNAi* largely failed to grow and did not show a significant increase in protein, as compared to controls (Figure 5A, B). This dramatic suppression of total gut protein was observed when *EcR-RNAi* was expressed under the control of either the *Tubulin-Gal4^ts^* (Figure 5A) or *Myo1A-Gal4^ts^*(Figure 5B) drivers, which are expressed in the whole animal (including gut) and specifically in enterocytes (ECs), respectively. Thus, the increase in the protein content of the gut after mating is EcR-dependent and is largely dependent on EcR function specifically in enterocytes. These results align nicely with previous reports that used progenitor-specific *esg-Gal4^ts^* driver ^14, 17^, showing that mating-dependent gut growth is EcR-dependent, with that additional insight that EcR signaling specifically in ECs plays a role in mating-dependent gut growth.

**Figure 5.**
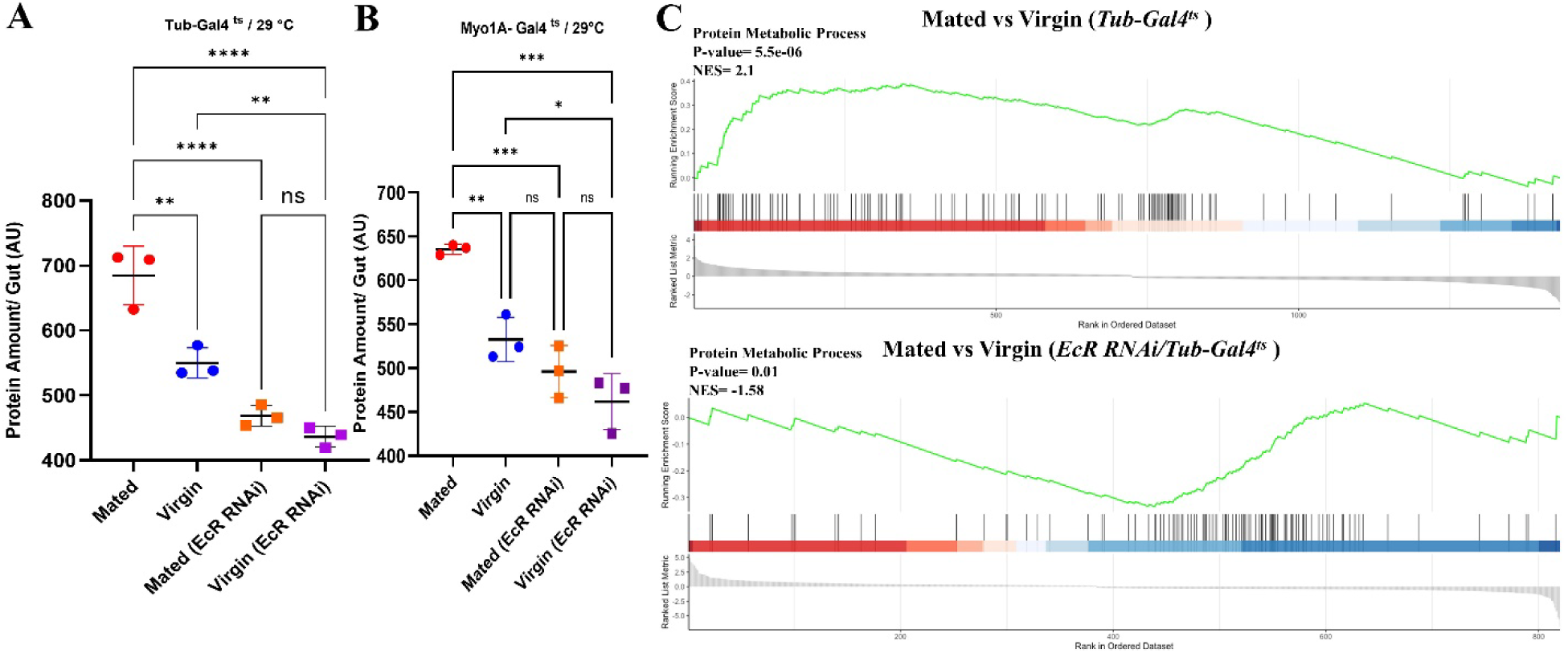
Mating-dependent upregulation of genes involved in protein metabolism requires EcR. **A-B.** Total protein content defined by BCA colorimetric assays comparing mated and virgin flies (each/3 replicates/15 guts) +/- *EcR- -RNAi* using *Tub-Gal4*^ts^ (A) and *Myo1A-Gal4^ts^* (B) drivers. **C.** GSEA comparing protein metabolic process genes expression in mated vs. virgin flies +/- *EcR-RNAi* using *Tubulin-Gal4*^ts^ driver. The Enrichment Score (ES) measures how much a gene set is overrepresented at either end of a ranking list, while the Normalized Enrichment Score (NES) reflects the degree of this enrichment, shown on the y-axis. GSEA normalizes the ES to account for differences in gene set sizes and their correlations with the expression data.

### Mating-dependent upregulation of protein metabolism genes are ecdysone signaling-dependent

To learn more about the regulation of mating-dependent increases in gut protein, we performed additional RNAseq analysis. We aged the collected virgin flies for 5 days at 29°C to express *EcR-RNAi* under the control of the *Tub-Gal4* driver. Then, after 5 days, we set the mating for 48 hours at 25°C by adding males, and we dissected the guts after 48 hours mating for RNAseq analysis. Mated females expressing *EcR-RNAi* under the control of the *Tubulin-Gal4^ts^* driver showed different transcriptome profiles compared to controls not expressing RNAi. Gene Ontology (GO) enrichment analysis of this data showed that GO terms related to protein metabolism were the most affected. Consistent with results shown earlier, GO terms such as proteolysis (GO: 0006508), amino acid transport (GO: 0006865), carboxylic acid transmembrane transport (GO: 1905039), and amino acid transmembrane transport (GO: 0003333) were significantly upregulated by mating in the midgut (Extended Figure 2B). However, the statistical significance of the upregulation of protein metabolism genes was massively affected in mated flies expressing *EcR-RNAi* under the control of the *Tub-Gal4^ts^* driver (Extended Figure 2B), suggesting that ecdysone signaling is critical for the regulation of these genes and processes after mating. Gene Set Enrichment Analysis (GSEA) of RNAseq data from mated flies expressing *EcR-RNAi* compared to mated controls confirmed that EcR signaling controls mating-induced changes in a broad range of protein metabolism genes, including genes involved in proteolysis and amino acid transport (Figure 5C). However, this EcR-specific effect was not observed when *EcR-RNAi* was expressed with the EC-specific *Myo1A-Gal4^ts^* driver (Extended Figure 2B, C), suggesting that EcR-mediated regulation of protein metabolism genes occurs in a non-EC cell type, possibly outside the midgut. This is consistent with recent studies indicating an important role for foregut cells in fly protein metabolism ^29^.

### Mating-dependent upregulation of *Jonah* protease genes requires Ecdysone/EcR signaling

Protein digestion requires both endo- and exo-peptidase enzymes. In *Drosophila*, *Jonah* family endopeptidases, *yip7* and *trypsin* first split large ingested proteins at internal peptide bonds, and the smaller peptides that result are in turn hydrolyzed to single amino acids, starting at their terminal ends, by exopeptidases such as carboxypeptidases (*CG12374*, *CG17633*), or to smaller peptides by di-peptidyl peptidases (*CG3744*, *CG11034*, *CG17684*, *Dpp10*, *ome*) ^30^. Subsequently, intestinal protein absorption requires amino acid transporters (*Slif, NAAT, path*) ^14, 31^ and oligopeptide transporters (*Yin*, *CG2930*, *CG9444*) ^31^ that transfer the digested protein into cells, most likely enterocytes. Our RNA-seq data showed that mating significantly increased the expression of many *Jonah* gene transcripts (*Jon25Bii, Jon65Aii, Jon99Cii, Jon99Ciii, Jon65Aiii, Jon65Aiv, Jon25Bi, Jon44E, Jon66Ci*, and *Jon66Cii*; Extended Figure 1A), which encode serine endo-peptidases used for digesting dietary protein. Mating also increased the expression of other peptidases (*trypsins*, *yip7*, *CG12374)*, as well as the amino acid importers *Slif*, *NAAT*, *dmGlut,* and *path*. As with the broader set of protein metabolism genes, *EcR-RNAi* under the control of *Tub-Gal4^ts^* significantly suppressed the mating-dependent upregulation of most of these genes (Extended Figure 2B). Surprisingly, this inhibitory effect was absent when *EcR-RNAi* was specifically expressed in enterocytes using the *Myo1A-Gal4^ts^* driver (Extended Figure 2B). Of note, the expression of protein absorption genes (*Yin*, *CG2930*, *CG9444*, *Slif*, *path*) was not affected by *EcR-RNAi* under the control of either driver. These results suggest that EcR may regulate the digestive steps of protein metabolism in mated females through a non-EC autonomous function (see also Data S4).

### Ecdysone signaling modulates both protein and lipid metabolism, with a stronger effect on protein metabolism in mated flies

Based on the observations above, we predicted that ecdysone signaling might preferentially regulate protein digestion and metabolism in mated females. Hence, we performed metabolite profiling on the guts of mated and virgin flies, each with or without *EcR-RNAi* conditionally expressed under the control of the *Tubulin-Gal4 (Tub-Gal4^ts^)* driver. We aged newly eclosed virgin flies for 5 days and shifted to 29°C to express *EcR-RNAi*. After 5 days, half the flies were mated for 48 hours at 25°C by adding males, whereas the rest (virgin samples) were cultured for 48 hours at 25°C without males. Guts were then dissected for metabolomics analyses. As noted above, the metabolites that changed the most after mating were fatty acids (linoleic acid, lauric acid) and TCA cycle intermediates (malic acid, fumaric acid), all of which were increased in the midguts of mated versus virgin females (Extended Figure 4A, Extended Figure 5A). However, in the midguts of mated flies expressing *EcR-RNAi,* these lipids, and TCA cycle intermediates were not significantly altered (Extended Figure 5B, 5C), indicating that mating-induced changes in those specific metabolites were disrupted by the loss of EcR. Instead, the metabolites with the most significant changes were L-serine and ornithine, which were both significantly decreased compared to virgin controls (Extended Figure 4A, Extended Figure 5C) (See also Data S1); these metabolites were not significantly affected by *EcR-RNAi* in virgin flies, showing that these effects are mating-specific (Extended Figure 5C) (See also Data S1). In addition to being required for protein synthesis, L-serine is an important metabolite in purine and pyrimidine synthesis, sphingosine, sphingolipid, and ceramide synthesis, and as a precursor for the biosynthesis of glycine, cysteine, tryptophan, and phospholipids. L-serine also supports the TCA cycle as an intermediate in the methionine cycle ^32^. Ornithine is involved in the urea cycle and can be used to generate intermediates of the TCA cycle, and it can be synthesized in the pyrimidine synthesis pathway. In summary, we can conclude that ecdysone signaling is a key regulator of diverse aspects of metabolism, specifically in mated flies.

Interestingly, our analyses showed that most changes in lipid metabolism were not affected by *EcR-RNAi*, regardless of mating status (Figures 4 A, B, C, 6A). Considering this and the effect of *EcR-RNAi* on ceramides and fatty acids enrichment in mated flies, as discussed above, we can conclude that EcR is partially involved in lipid metabolism in mated flies. However, changes in metabolites associated with protein metabolism, such as changes in amino acids, were largely EcR-dependent in mated flies (Figure 6B; see also Data S1). Hence, we surmise that EcR signaling controls both protein and lipid metabolism but has a stronger influence on protein metabolism. In conclusion, loss of EcR altered the metabolic profile of mated flies by interfering with metabolic pathways responsible for the provision of TCA intermediates, amino acids, ceramides, and acid nucleic synthesis. These metabolic effects may be important for the ecdysone signaling-mediated activation of mating-dependent ISC proliferation and growth in the midgut (Figures 1, 5A) ^17, 18^.

**Figure 6.**
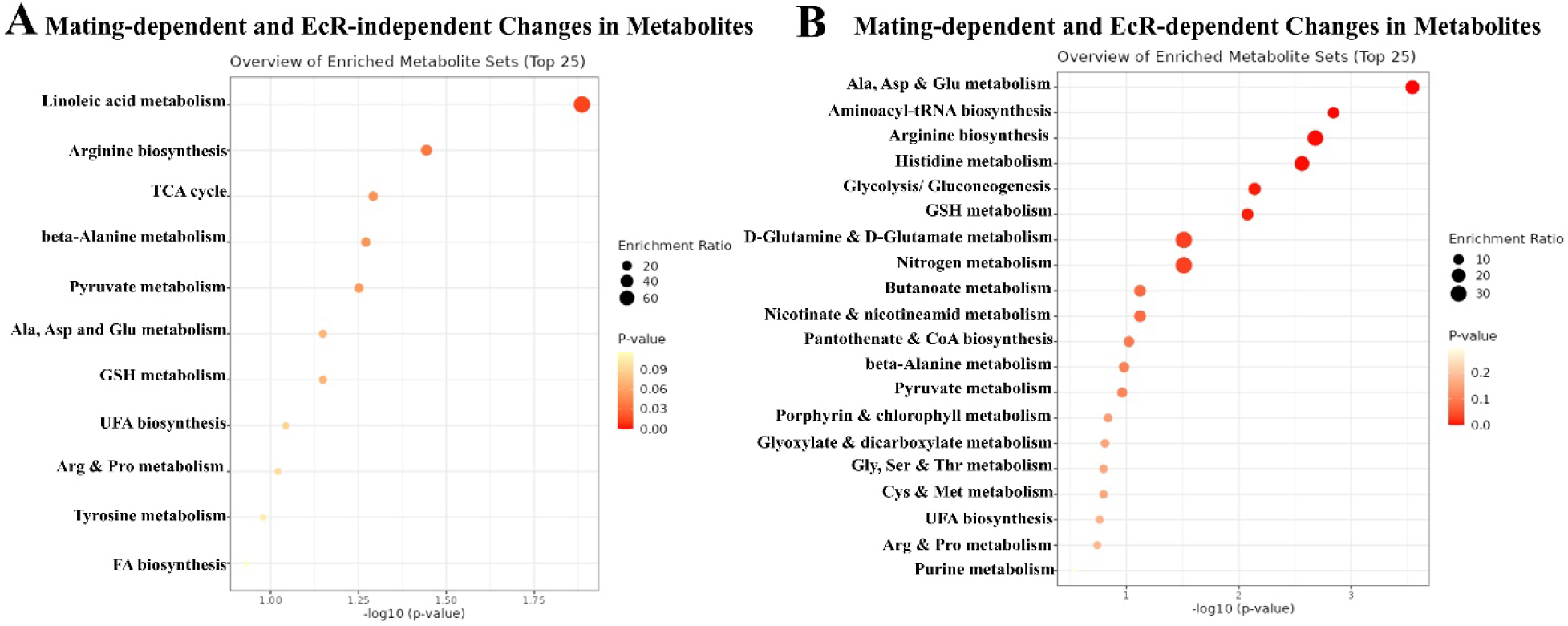
EcR significantly affects metabolic processes related to protein metabolism. **A-B.** GC-MS Metabolite set enrichment analysis of changes in metabolites comparing mated and virgin flies +/- *EcR-RNAi* using *Tub-Gal4ts* (each/4 replicates/ 60 guts) showing changes in metabolic processes upon mating, the analysis revealed changes in metabolic processes triggered by mating that occur independently of EcR expression (A), as well as those specific to mated flies without *EcR-RNAi* knockdown (B).

### Mating Promotes Protein Digestion Efficiency

Several studies have emphasized the importance of dietary protein ^33^ and protein metabolism ^34, 35^ for egg production by *Drosophila* females. Based on the observations detailed above (*e.g.,* Figure. 3), we postulated that not only food preference ^8, 9, 13, 14, 36^, but also digestive function might be altered by mating. Hence, we assessed the efficiency of protein and carbohydrate digestion in mated females by performing colorimetric assays of the protein and carbohydrate contents of their feces (see Methods). Standard fly media was used in all cases. We found that mated females, despite having a larger gut (Figure 1) and eating more than twice as much as virgins ^10, 37^, excreted only a small amount more total protein in their feces than virgins (∼9 vs 6 AU; Figure 7A). However, in line with their increased food consumption, mated flies did excrete much more carbohydrates in their feces than virgins (20 vs 4 AU; Figure 7A). When viewed as protein/carbohydrate ratios (Figure 7B), this data clearly indicates more efficient relative protein digestion after mating. Notably, however, depleting the ecdysone receptor (EcR) in enterocytes (genotype: *MyoIA-Gal4^ts^>UAS-EcR^RNAi^*) did not significantly change the amounts or ratios of protein and carbohydrate in the feces (Figure 7B). Given that we expect the enzymes and transporters that mediate digestion to act in and be expressed by enterocytes, this is somewhat surprising. As a potential explanation, we note that while our RNAseq experiments showed that the expression of many protein and carbohydrate digestive enzymes and transporters was significantly altered by mating (*e.g.,* protease expression was increased and maltase expression decreased; Figure 3C, D), these effects were not appreciably suppressed by *EcR-RNAi* driven under the control of the EC-specific *MyoIA-Gal4* driver (Extended Figure 2B, C). On the other hand, we did see a significant suppression of the mating-dependent induction of protein digestion genes when *EcR-RNAi* was expressed under the control of the ubiquitous Gal4 driver, *Tubulin-Gal4^ts^* (Figure 5C, Extended Figure 2A, B). However, in this case, carbohydrate digestive enzymes were still not highly affected (Extended Figure 3B, C). Similarly, mating-induced transcriptional downregulation of autophagy-related genes was not affected by *EcR RNAi* (Extended Figure 3D). These observations suggest that there probably are EcR-dependent genes that are needed for the switch to more efficient protein digestion upon mating, but that some of these genes are expressed in a cell type other than enterocytes, and which might not even reside in the midgut. This non-EC autonomous effect of the *EcR-RNAi* might be explained, for instance, by a signaling factor that is induced upon mating by ecdysone/EcR in the brain or fat body, and which indirectly regulates *Jonah* protease gene expression in midgut enterocytes. In summary, these results support the idea that more efficient protein digestion is mating-dependent but not directly dependent upon EcR function in enterocytes.

**Figure 7.**
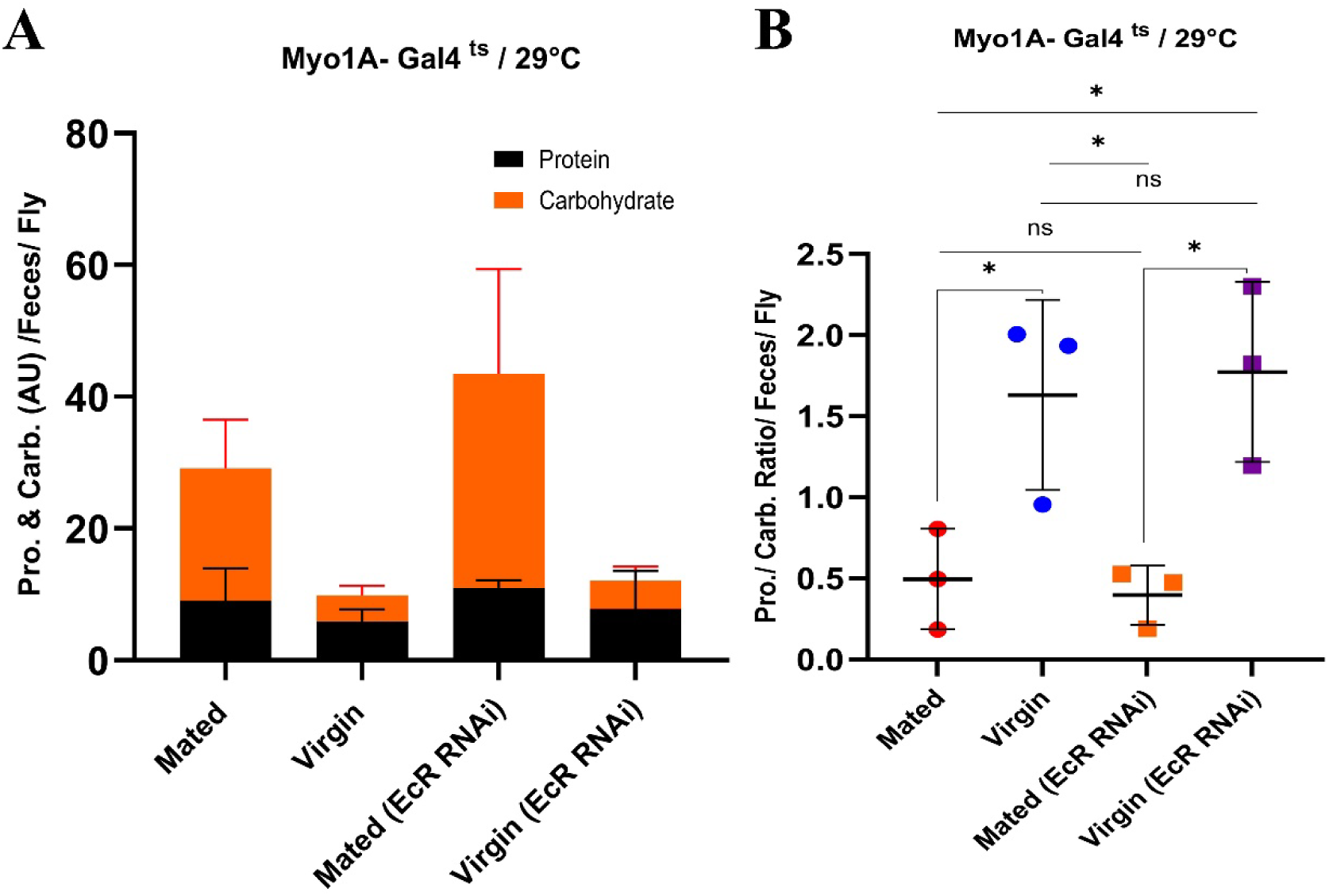
Mating enhances protein digestion efficiency. **A.** Proportion of carbohydrate and protein in feces of mated and virgin flies +/- *EcR-RNAi* (*Myo1A-Gal4^ts^*). **B.** Protein/Carbohydrate ratios in mated and virgin flies +/- *EcR-RNAi* (*Myo1A-Gal4^ts^*).

## DISCUSSION

When we consider normal intestinal physiology, it is notable that adaptive growth of the intestine is well documented in pregnant and lactating female mammals of several species ^38-40^. The onset of this intestinal growth coincides with the massive increases in estrogen, progesterone, and corticol steroids that occur during gestation. Estrogen, progesterone, and cortisol levels drop precipitously postpartum, but intestinal growth continues during lactation when levels of the prolactin hormone remain high. Like estrogen and progesterone ^41^, ecdysone is a cholesterol-derived female-biased steroid hormone that controls aspects of sexual behavior, dimorphism, physiology, and reproductive function in insects ^42^. The similarity in the structures of these hormones, their sites of synthesis, regulated secretion, and actions on the reproductive system suggests that they could have similar actions in other organs, too. If increased estrogen or progesterone titers in pregnant mammals are causal for intestinal growth (this has not yet been rigorously tested), their functions in mammals would be analogous to the effects of ecdysone in mated *Drosophila* females ^17, 18^. Previous reports, however, did not address how adaptive gut growth after mating is accompanied by different metabolic adaptations and whether the effects are steroid signaling-dependent. We report on these topics in *Drosophila* here.

We report that mating induces significant physiological and metabolic changes in the female *Drosophila* gut. Consistent with previous studies, we observed adaptive growth of the gut mediated by ecdysone signaling ^17, 18^. This expansion is characterized by a larger gut size, more total protein per gut, and more total cells, including progenitor cells. Mating-induced intestinal growth is also accompanied by major changes in the gut’s biochemical composition. We observed large, significant increases in total gut lipids and protein per gut in mated flies, while carbohydrate amounts were relatively unchanged. Our metabolomics analyses revealed that several TCA cycle intermediates increased significantly in mated female guts, including fumaric and malic acids. TCA cycle products, including NADH and FADH2, power the mitochondrial electron transport chain and the synthesis of ATP ^43^, and so the changes we observed likely reflect alterations in ATP utilization. Fumarate also acts as an electrophile and induces the succination of proteins, whereby fumarate binds to reactive thiol groups in cysteine residues of proteins and inactivates them. Fumarate can also reshape the epigenetic landscape by inhibiting histone and DNA demethylases ^43, 23^. Through these two avenues, altered Fumarate levels may have widespread effects on gene expression. Another key metabolite that significantly increased in the gut of mated flies is myo-inositol, a precursor of the essential membrane phospholipid, phosphatidylinositol (PI), which has roles in signal transduction, endoplasmic reticulum stress (unfolded protein response), energy metabolism, nucleic acid synthesis, and osmoregulation ^44^. These examples from our metabolite profiling experiments highlight that mating not only increases the total amounts of macromolecules, including proteins and lipids per gut but affects myriad details of a wide spectrum of metabolic pathways, all presumably to maximize biosynthesis and energy production for the generation of large quantities of eggs, which are relatively massive.

LC-MS analysis also showed an overall increase in lipid content in the guts of mated flies that was even greater than the increase in protein content. This may reflect a greater demand for stored energy in the form of lipids after mating. Triglycerides (TG) and ceramides were amongst the top classes of lipids with significant changes. As an important storage form of lipids, TGs are important for sustained energy production. Ceramides are not only involved in membrane structure but are also involved in signal transduction and enhance the proliferation of progenitor cells in the gut ^28^. Metabolomics analyses also confirmed increases in fatty acids, which are used as precursors to many of the more complex lipids used in cell growth. In addition, fatty acids can be an important alternative energy source to fuel the TCA cycle and ATP production. Furthermore, the lipid changes induced by mating, particularly increased ceramides, appeared to be partially dependent on ecdysone signaling. A previous study indicated that mating changes lipid metabolism through the activation of Sterol Regulatory Element Binding Protein (SREBP) ^19^; however, this gene was not detected as transcriptionally upregulated in our RNAseq data sets. Nevertheless, the overall increase in total lipid content, including key fatty acids such as linoleic and lauric acids, supports enhanced energy storage and utilization during reproduction, as previously discussed ^12, 19^.

Interestingly, gene expression profiling showed that mating significantly upregulated *Jonah* family peptidase genes and other genes involved in protein digestion, such as the amino acid and small peptide transporters required for the intestine to absorb proteolyzed proteins. Conversely, genes used in carbohydrate metabolism and digestion were relatively downregulated. This is consistent with our observation that mating caused a relative reduction in gut carbohydrate content and a large increase in the carbohydrate content of the feces. In agreement with our results, White et al. ^14^, Gioti et al.^45^, and Zhou et al. ^46^ previously detected a similar post-mating up-regulation of *Jonah* endopeptidase genes. Reiff et al. ^19^ used qPCR to detect the up-regulation of several fatty acid metabolic genes in the midgut after mating, and White et al. ^14^ noted the transcriptional upregulation of amino acid transporters, including *NAAT* and *Slif*. Our findings show additionally that aspects of the mating-induced switch towards protein digestion are EcR-dependent. This underscores the idea that ecdysone-dependent protein metabolism in female flies is prioritized after mating and is likely to meet the increased biosynthetic demands of reproduction. Hudry *et al.* ^26, 47^ showed that genes used in cell division-related processes are more abundantly expressed in females, whereas genes encoding functions in carbohydrate metabolism and redox processes are preferentially expressed in males. This and other observations suggest physiological similarities between virgin females and males, and imply that the mating-induced switch towards protein metabolism, promoted by higher ecdysone titers ^48^, could be a female-biased phenomenon.

The upregulation of protein metabolism genes is unlikely to be merely a consequence of increased food intake since we also observed simultaneous downregulation of carbohydrate metabolism genes. Instead, we propose that the female gut changes its digestive parameters after mating to adapt to the new nutritional requirements mandated by egg production. A previous study by White et al. ^14^ also suggested this possibility, whereas others mostly focused on the fly’s post-mating dietary preference for a protein-rich diet^8-11, 13, 33, 34^.

Our transcriptome profiling also revealed a significant, widespread mating-dependent down-regulation of genes involved in stress responses, especially oxidative stress-associated genes. This effect was far stronger in mRNA from *esg^+^* intestinal stem cells (ISC) and enteroblasts than in whole midgut mRNA. This intriguing observation suggests that ISCs in mated flies may have a unique ability to cope with Reactive Oxygen Species (ROS), which are produced by mitochondria and the gut microbiota and are expected to rise in mated females’ nutritional throughput and energy production increase. We suggest that this phenomenon may be due to the increased protein metabolism in mated females, which leads to increased levels of methionine, a precursor of glutathione, an important ROS neutralizer ^49^. Increased glutathione might enable mated flies to cope with oxidative stress using the basal-level expression of *glutathione-S-transferase* (*GST)* genes.

Our RNAseq data from whole-gut mRNA also revealed a widespread downregulation of autophagy-related genes upon mating. Autophagic activity is typically low under growth-promoting conditions, such as during gut growth and egg production in our study, and is conversely elevated under growth-limiting conditions, such as during metabolic stress or nutrient deprivation ^50^. Our finding that autophagy genes were downregulated upon mating is consistent with a report that mutants in the autophagy gene *Atg101* have a thicker midgut epithelium with enlarged enterocytes ^51^ and suggests that the suppression of autophagy may aid enterocyte growth and/or digestive functions after mating. Interestingly, our data also showed that autophagy gene expression was not generally downregulated in *esg+* progenitor cells after mating, where, in fact, many autophagy genes were induced (Extended Figure 3A). These observations suggest different roles and modes of regulation of autophagy in ISCs and enterocytes and parallel reports that mTOR activity, a suppressor of autophagy, rises during ISC to enterocyte differentiation^52, 53^. Genetic tests of the requirement for autophagy in ISCs have given contradictory results, with one report showing ISC loss from ATG gene suppression^54^ and another documenting increased ISC proliferation^55^.

While food intake increases after mating, the intestinal transit rate of the food is significantly decreased. This has been shown to be due to the action of a male-derived reproductive hormone, Sex Peptide, which concentrates the intestinal contents and slows intestinal food transit in mated females ^56, 57^. This mating-dependent gastrointestinal change may be beneficial as a means to increase nutrient absorption at a nutritionally demanding stage for the female ^56^. Our results show that mating does indeed increase protein absorption, as evidenced by a relative reduction in protein content in the feces of mated females. This increase in protein digestion efficiency appeared not to require the Ecdysone Receptor (EcR) in enterocytes, despite our observation that ecdysone signaling somewhere in the fly was necessary for full induction of a large set of genes in the gut that encode digestive peptidases and peptide and amino acid importers. Although further experiments are required to resolve the mechanism, our data as they stand are consistent with the possibility that the switch to increased efficiency of protein digestion is mediated by the induction of peptidases and importers in the midgut but that EcR controls this induction indirectly, acting in a cell type other than enterocytes, perhaps located in another organ. This would implicate the involvement of a secondary signaling system as a relay. Alternatively, our observations could be explained by the existence of key genes that are directly used in protein digestion and which are directly controlled by EcR but in a gut cell type other than enterocytes, for instance, enteroendocrine or foregut cells ^29^. Finally, it is also possible that the switch to enhanced protein digestive efficiency is not regulated by ecdysone signaling at all but is regulated by another mating-dependent signal, such as Sex Peptide ^14, 45, 57^, Juvenile Hormone^19^, or Neuropeptide F ^13^.

Cognigni *et al*. ^56^ reported that *Drosophila* fecal deposits are acidic when flies are fed a sucrose diet but not when they are fed a high-protein diet. They also found that reproduction affects the acid-base balance of the gut, such that mated, egg-producing females showed similar acidification of excreta (feces) as flies cultured on high sugar/low protein diets ^56^. Based on these observations, Cognigni *et al. ^56^* proposed that the fly midgut “can modulate the final composition of intestinal contents” (*i.e.,* the feces) and that the metabolic demands of reproduction in female control an important modulatory input. Our results are consistent with this pattern, as we found that mated flies have a higher ratio of carbohydrate to protein in their feces. Interestingly, older work on mammals, summarized by Charney and Dagher ^58^, references similar changes in acid/base balance that also suggested the gut’s ability to modulate its digestion functions. Our analyses, showing strongly altered carbohydrate/protein balance in the feces following mating, align nicely with the results of Cognigni et al. ^56^ and add more direct evidence for the ability of the gut to rapidly adapt to extra-intestinal stimuli (*e.g.,* mating) by altering its digestive functions through changes in digestive gene expression and altered metabolic controls.

In summary, mating in female *Drosophila melanogaster* triggers extensive physiological and metabolic adaptations in the gut that optimize protein digestion and alter lipid metabolism. We presume that these changes optimize nutrient intake in order to support reproduction. These results advance our understanding of the intricate interplay between mating, gut physiology, and metabolic regulation and highlight the dynamic nature of biomolecular responses to reproductive stimuli. Future research into the molecular mechanisms underlying these adaptations and their impact on reproductive fitness and metabolic health should be very informative. Our analysis of the interplay between gene expression and metabolomics had clear limitations, as we performed only steady-state metabolite profiling and did not do nutrient uptake or metabolite flux assays. Both approaches could give even more detailed insight into how gut physiology and metabolism are altered by mating.

We hope our findings will help to stimulate and guide investigations of how mammalian sex hormones and reproduction in general, affect various aspects of physiology, metabolism, and digestion in the intestine. Mapping steroid- and reproduction-dependent changes in gut physiology and metabolism can shed light on the risks associated with adaptive gut growth during pregnancy, lactation, and/or puberty. Tracking sex hormone-mediated changes in gut metabolism in mammals may also be useful to evaluate the physiological effects and health risks of steroids used clinically, for instance, in hormone replacement and cancer therapies and sports doping, and to identify new ways to diagnose sex hormone-mediated metabolic disorders.

## MATERIAL AND METHODS

### Drosophila stocks and husbandry

*Drosophila melanogaster* flies were grown on standard media and housed in incubators under controlled temperature and humidity conditions in a 12-hour light-dark cycle. Culture vials containing the flies were replaced with fresh vials every 2 days. Control groups were generated by crossing *w^1118^*(VDRC #60000) flies with the corresponding Gal4 driver line. For transgene expression using the *Gal4/Gal80^ts^* system, experimental crosses were maintained at 18°C (permissive temperature for *GAL80^ts^*) in a standard medium. Animals of the desired sex and genotype were collected on Day 0 after eclosion and aged for 5 days at 29°C (restrictive temperature for *GAL80*^ts^) to induce UAS transgene expression. Adult guts were dissected after 48 h mating, as explained below, in the “mating experiment.”

### Mating experiments

At least 10–15 virgin females of each genotype, for each condition and replicate, were collected at 18°C upon eclosion. They were then aged at 25°C for 4-5 days or at 29°C (in experiments using *EcR-RNAi*) for approximately 5 days until mating began. At the onset of mating, virgin females were transferred to new vials with food and mated at 25°C for 48 hours with an equivalent number of adult wild-type *w^1118^* males that were between 3–7 days old at 25°C to maximize fecundity. After 48 hours of mating, the males were discarded, and the females were used for gut dissection.

### Drosophila stocks and transgenes

*esg-Gal4, UAS-GFP/CyO; Tub-Gal80^ts^/TM6B* (Edgar Lab stock collections /University of Utah, U.S.A.).

*Myo1A-Gal4, Tub-Gal80^ts^/CyO* (Edgar Lab stock collections /University of Utah, U.S.A.).

*Tub-Gal80^ts^/CyO* (Edgar Lab stock collections /University of Utah, U.S.A.).

*y[1],v[1];P{y[+t7.7]v[+t1.8]=TRiP.JF02538attP2* (EcR JF02538) ( Bloomington *Drosophila* Stock Center, BL29374).

### Flow Cytometry

To improve the efficiency of nuclear isolation for sorting nuclei by flow cytometry, we used a refined mating protocol that differs from the mating configuration used for the remainder of this study, as described previously. Virgin *Drosophila melanogaster* flies at 5 days post-eclosion age (maintained at 25°C) were selected for the mating experiment. A cohort of 25 flies was mated with a corresponding number of *w^1118^* males over a period of 5 days at 25°C, in addition to 25 age- and genotype-matched virgin control flies kept in a separate vial. After the five-day mating period (10 days post-eclosion), flies gut were carefully dissected for nuclei isolation.

For both conditions, a Nuclear Extraction Buffer (NEB) was prepared by mixing 715 µL Pre-Nuclear Extraction Buffer, 35.75 µL 25X Protease Inhibitor, 0.715 µL DAPI dye and 0.357 µL 1 M spermidine. Prior to dissection, pipette tips and tubes were coated with 10% Normal Goat Serum (NGS) in 1% PBTX. The Drosophila midguts were dissected in chilled 1x phosphate-buffered saline (PBS) and then transferred to a cold glass vial coated with 10% NGS and filled with 250 µL NEB. In the well, the midguts were crushed into small pieces using forceps. The samples were then placed in a 1.5 mL low-binding tube and incubated on ice in a Nutator for 15 minutes. After incubation, samples were centrifuged at 4°C and 6000 rpm for 3 minutes and then resuspended in 400 µL NEB. Samples were gently mixed by pipetting for 20-30 cycles. To remove large debris, samples were filtered through 70-µm filters into new 10% NGS-coated 1.5-mL low-binding tubes. After filtering, samples were fixed to prevent degradation of the nucleus by adding 2.5 µL of 16% paraformaldehyde (PFA) for a final concentration of 0.1%. The samples were gently shaken and fixed at room temperature for 2 minutes. To neutralize the PFA, 12 µL of 2.5 M glycine was added, and the samples were briefly shaken again before centrifugation at 4°C and 1300 x g for 4 minutes. After centrifugation, the PFA supernatant was removed, and the samples were resuspended in 400 µL NEB with 20-30 slow pipetting cycles. Once core isolation was complete, samples were transferred to an Aria Cell Sorter on ice. With the help of the Huntsman Cancer Institute Flow Cytometry Core technicians, the nuclei were gated and sorted. The cytometer processed the samples at a rate of 2.5 µL per second, capturing all 400 µL of NEB and nuclei.

The data from the cell sorter was then analyzed in FlowJo using a six-step gating scheme to remove debris and isolate nuclei. The gating strategy included using different forward and side scatter channels to eliminate small, non-spherical, and elongated debris, gating non-DAPI+ nuclear populations, and separating nuclei by ploidy levels from 2C to 64C. See details below:

1. Set the SSC-A and FSC-A gates to identify the nuclei (based on the histogram plot) as determined by size and granularity (gate P1).
2. Set the FSC-A and FSC-H gates to identify nuclei singlets based on their size (gate P2).
3. Set the SSC-A and SSC-H gates to identify nuclei singlets based on their granularity (gate P3).
4. Set the SSC-A and DAPI Intensity gates to identify DAPI-positive Nuclei (gate P4).
5. Use gate P4 to depict the DAPI-positive cells in a histogram plot that shows the number of cells (count) vs. DAPI intensity. 6 distinct peaks of DAPI-positive cells will be visible. Create a gate for each peak. Each new gate contains the cell population that is enriched for cell ploidy from 2 C-64 C and can be sorted separately from each other.

After gating, the data sets were visualized as histograms in FlowJo’s layout editor, with DAPI fluorescence (BV421-A) on the x-axis and the number of nuclei on the y-axis. The data was then exported to GraphPad Prism to create the ploidy distribution and proportion of the whole plots.

### RNA sequencing

RNA isolation and amplification from sorted cells and whole midguts were performed as previously described ^59^. Four independent biological replicates for each condition were used for sequencing.

### Colorimetric Protein and Carbohydrate Assays

Samples for measuring protein and carbohydrate concentration in the Drosophila midgut were prepared by dissecting 10 to 15 midguts in 1X PBS. The carbohydrate concentration of this sample was measured using the Total Carbohydrate Quantification Assay Kit (Abcam, #ab155891) as per the manufacturer’s instructions. For measuring protein concentration, samples of 10-15 midguts were further processed by adding 200 µL of RIPA buffer and 4 µL of protease inhibitor. The midgut structure was broken up and homogenized by passing samples through a 26-gauge needle 10-15 times. Samples were left to sit on ice for 10 minutes, then centrifuged at 4°C and 13,000 rpm for 10 minutes. Protein concentration in the resulting supernatant was measured using the Pierce BCA protein assay kit (Thermo Scientific, #23227) as per the manufacturer’s instructions.

Samples for measuring protein and carbohydrate concentration in Drosophila feces were prepared by adding 50-70 flies to a wide vial with approximately 1 inch of lab-made food. Flies were kept for 24 hours before being discarded. Feces samples were collected by scraping the inside of the vial with a cotton swab damp with MiliQ water, taking care to avoid the food. Sample was suspended in 300-500 µL MiliQ water. Half of the sample was used to measure carbohydrates using the Total Carbohydrate Quantification Assay Kit (abcam, #ab155891), and half was used to measure protein. For measuring protein concentration, samples were further processed by adding RIPA buffer to feces solution (1:1) and protease inhibitor (1:200). Samples were set on ice for 10 minutes and then centrifuged at 4°C and 13,000 rpm for 10 minutes. Protein concentration in the resulting supernatant was measured using the Pierce BCA protein assay kit (Thermo Scientific, #23227) as per the manufacturer’s instructions.

RIPA Buffer (following recipe):

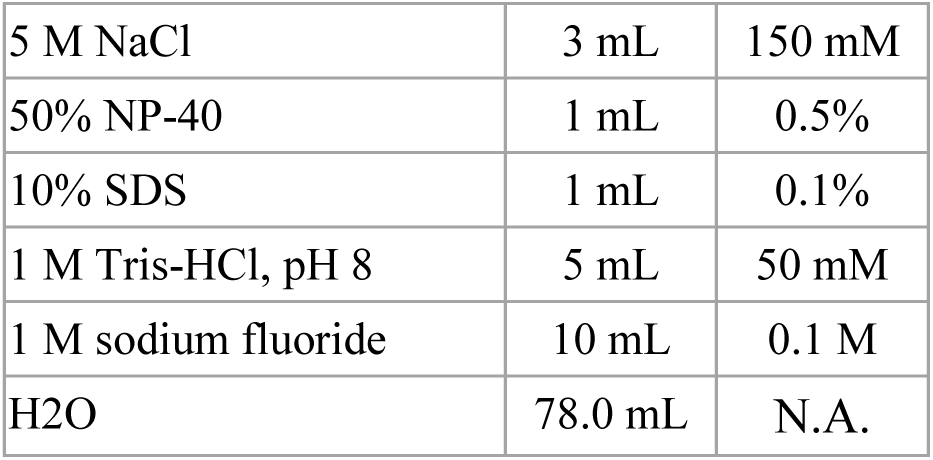

### Metabolomics and Lipidomics

The flies were dissected in ice-cold phosphate-buffered saline (PBS), and then their intestinal tissue was rapidly cryopreserved in liquid nitrogen. From each series of experiments, 60 and 40 intestines were collected for subsequent metabolomics and lipidomics analysis, respectively. Extraction of metabolites was performed by transferring each biological sample to 2.0 mL ceramic beadmill tubes (Qiagen catalog number 13116-50) and adding 450 mL of chilled 90% methanol (MeOH) solution containing the internal standard d4 succinic acid (Sigma 293075). Using an OMNI Bead Ruptor 24, the samples were homogenized and then incubated at -20 °C for one hour. After centrifugation at 20,000 x g for 10 minutes at 4 °C, 400 mL of supernatant from each tube was carefully transferred to appropriately labeled microcentrifuge tubes to which an additional internal standard, d27-myristic acid, was added. Quality control samples were carefully prepared by pooling a segment of the collected supernatant from each individual sample, while control blanks consisting only of the extraction solvent were subjected to the same processing protocols as the experimental samples. Subsequently, all samples were dried under vacuum conditions. Gas chromatography-mass spectrometry (GC-MS) analysis was performed using an Agilent 7200 GC-QTOF equipped with an Agilent 7693A automated liquid collector. The dried samples were reconstituted in 40 mL of a 40 mg/mL O-methoxylamine hydrochloride (MOX) solution dissolved in dry pyridine and incubated for one hour at 37 °C in a sand bath. After this incubation time, 25 mL of the resulting solution was transferred to autosampler vials to which 60 mL of N-methyl-N-trimethylsilyltrifluoroacetamide (MSTFA with 1% TMCS, Thermo #TS48913) was automatically added via the autosampler and then incubated at 37 °C for 30 minutes. After incubation, the samples were shaken, and 1 mL of the prepared sample was injected into the gas chromatograph in split mode at an inlet temperature of 250 °C. The analysis was performed with a split ratio of 10:1 for most metabolites. For those metabolites that saturated the device at a split ratio of 10:1, the split ratio was changed to 50:1. Chromatographic separation was performed with a 30 m Agilent Zorbax DB-5MS coupled to a 10 m Duraguard capillary column using helium as the carrier gas at a flow rate of 1 mL/min.

The method of lipid extraction was based on the protocols of Matyash et al. (J Lipid Res 49(5) (2008) 1137-1146). Lipid extracts were separated using an Acquity UPLC CSH C18 column (2.1 x 100 mm; 1.7 mm) in conjunction with an Acquity UPLC CSH C18 VanGuard precolumn (5 x 2.1 mm; 1.7 mm) (Waters, Milford, MA). The operating parameters of this system were controlled to a temperature of 65°C and were connected to an Agilent HiP 1290 sampler, an Agilent 1290 Infinity pump, and an Agilent 6545 Accurate Mass Q-TOF dual AJS-ESI mass spectrometer (Agilent Technologies, Santa Clara, CA). Samples were systematically analyzed in a randomized sequence using both positive and negative ionization modes over a scan range of m/z 100 – 1700. In the positive ionization mode, the source gas temperature was calibrated to 225°C, accompanied by a drying gas flow rate of 11 l/min, an atomizer pressure of 40 psig, a sheath gas temperature of 350°C, and a sheath gas flow rate of 11 l/min. Conversely, for the negative ionization mode, the source gas temperature was set to 300°C, with a drying gas flow rate of 11 L/min, an atomizer pressure of 30 psig, a sheath gas temperature of 350°C, and a sheath gas flow rate of 11 L/min. Mobile phase A consisted of ACN:H2O (60:40 v/v) in 10 mM ammonium formate with 0.1% formic acid, while mobile phase B was formulated as IPA:ACN:H2O (90:9:1 v/v/v) in 10 mM ammonium formate together with 0.1% formic acid. For the analysis performed in negative mode, the modifiers were replaced by 10 mM ammonium acetate. The chromatographic gradient for both the positive and negative ionization modes started at 15% B of the mobile phase and increased to 30% B over a period of 2.4 minutes, followed by an increase to 48% B from 2.4 to 3.0 minutes, a further increase to 82% B from 3 to 13.2 minutes, a subsequent increase to 99% B from 13.2 to 13.8 minutes, which was then maintained until 16.7 minutes and finally returned to the initial conditions, with a 5-minute equilibration period. The flow rate was kept constant at 0.4 mL/min, with injection volumes set at 1 mL for the positive ionization mode and 10 mL for the negative ionization mode. Tandem mass spectrometry was performed with iterative exclusion with collision energies of 20 V and 27.5 V in positive and negative modes, respectively.

### mRNA sequence analysis

The sample reads were aligned to the latest genome assembly (BDGP 6.94) using the STAR Aligner. The aligned reads were then assigned to genes using HTSeq in ‘Union’ mode. The raw scores derived from this process were then used for differential expression analysis using the DESeq2 algorithm. Genes that met the criteria of an absolute log2 fold change ≥1 and an adjusted p-value <0.05 were identified as significantly differentially expressed.

### Gene Ontology enrichment analysis

Gene Ontology biological process terms enrichment analyses were conducted using the R package topGO68 and the Drosophila melanogaster annotation file released on 2019-07-01 from the Gene Ontology Consortium website.

### Heatmaps

Heatmaps were generated using the heatmap.2 function from the R package gplots (https://cran.r-project.org/package=gplots). Raw reads were converted to Transcripts Per Million (TPM), z-score normalized, manually grouped and ordered, and plotted without unsupervised clustering.

### Gene Set Enrichement Analysis (GSEA)

(http://www.gsea-msigdb.org/) Genes from RNASeq data were ranked according to their log2 fold-change and tested for enrichment of specific metabolic gene sets downloaded from the KEGG database. The most important result of Gene Set Enrichment Analysis is the Enrichment Score (ES), which indicates the extent to which a gene set is disproportionately represented either at the top or bottom end of a ranking list. The Gene Set Enrichment Analysis (GSEA) calculates the ES by running through the ranked gene list and increasing a running sum statistic when a gene is included in the gene set and decreasing it when it is excluded. The extent of the increase depends on the correlation between the gene and the phenotype under study. The ES represents the highest deviation from zero observed during the run of the list. A positive ES means an enrichment of genes in the upper part of the ranking list, while a negative ES means an enrichment of genes in the lower segment. In graphical representations of the GSEA, the normalized enrichment score (NES) reflects the degree of enrichment and is shown along the y-axis. The NES values serve as the basis for ranking the gene sets on the x-axis. The NES is derived by normalizing both the positive and negative enrichment scores (ES) against the mean of the respective positive or negative pES. This adjustment accounts for variations in the size of the gene sets as well as correlations between the gene sets and the expression data set. The normalized enrichment score (NES) is the main statistic used to evaluate the results of gene set enrichment analyses. By normalizing the enrichment score, GSEA compensates for differences in gene set size and correlations between the gene sets and the expression data set. Consequently, normalized enrichment scores (NES) facilitate comparative analyses of results across multiple gene sets.

### LC-MS data processing

A comprehensive methodology was applied for data processing, including the use of the Agilent MassHunter (MH) workstation and the MH Qualitative and MH Quantitative software packages. To ensure the reliability of our lipidomics dataset, pooled quality control samples (n=8) and blanks (n=4) were systematically integrated into the sample list. Lipid annotation was performed with great care, using precise mass and MS/MS correspondence using the Agilent Lipid Annotator library and LipidMatch (as described below). Lipid Annotator-derived results for both positive and negative ionization modalities were merged based on classified lipid identification. Subsequently, the data extracted from M.H. Quantitative was subjected to a comprehensive review in Excel, where the initial lipid targets were examined according to established criteria. Lipids with relative standard deviations (RSD) of less than 30% within the QC samples were prioritized for further investigation. In addition, lipids with background AUC values in blank samples that were less than 30% of those found in the QC samples were included in the analysis. The organized Excel data tables were then normalized using class-specific internal standards, followed by normalization to tissue mass and aggregation prior to subsequent statistical analysis.

### Metabolomics and lipidomics data analysis

The peak data was carefully analyzed using the MetaboAnalyst software platform. The log2-fold changes were calculated on the basis of the unprocessed peak values. Before performing the statistical analysis, the raw peak data were subjected to a log10 transformation. Then, the peaks corresponding to each metabolite were standardized by centering them on the mean and normalizing them by the square root of their respective standard deviation, a technique commonly referred to as Pareto scaling.

## ACKNOWLEDGEMENTS

This work was supported by National Institute of Health Grant R01 DK125745 (to B.A.E). We thank the Bloomington Drosophila Stock Center for Drosophila. We thank the Huntsman Cancer Institute High-Throughput Genomics and Bioinformatic Analysis Shared Resources, the University of Utah 3i-UCGD Bioinformatics Core, and the University of Utah Metabolomics and Flow Cytometry cores for assistance. Our Metabolomics Core was supported by NCRR shared instrumentation grants 1S10OD016232-01, 1S10OD018210-01A1, and 1S10OD021505-01. Our Flow Cytometry Core was supported by N.I.H. director’s office award S10OD026959 and NCI award 5P30CA042014-24.

## SUPPLEMENTARY INFORMATION

**Data S1.** Metabolomics lipidomics data of mated and virgin flies, related to Figure 2, Figure 4, Figure 6, Extended Data Figure 4, and Extended Data Figure 5.

**Data S2.** Whole gut and FACS-sorted esg+ mRNA-seq summary tables data of mated and virgin flies, related to Figure 3, Extended Figure 3A.

**Data S3.** The full list of enriched gene ontology biological processes in terms of mated vs virgin up- or down-regulated genes is related to Figure 3A-B.

**Data S4.** mRNA-seq summary tables data of mated and virgin flies +/- *EcR-RNAi* under the control of *Tub-Gal4^ts^* and *Myo1A-Gal4^ts^* related to Figure 5C, Extended Figure 2 and Extended Data Figure 3 B-D.

**Data S5.** Full list of enriched gene ontology biological processes terms of mated *vs* virgin +/- *EcR-RNAi* under the control of *Tub-Gal4^ts^* and *Myo1A-Gal4^ts^*, up-regulated genes, related to Extended Figure 2B.

## EXTENDED DATA

**Extended Data Figure 1.**
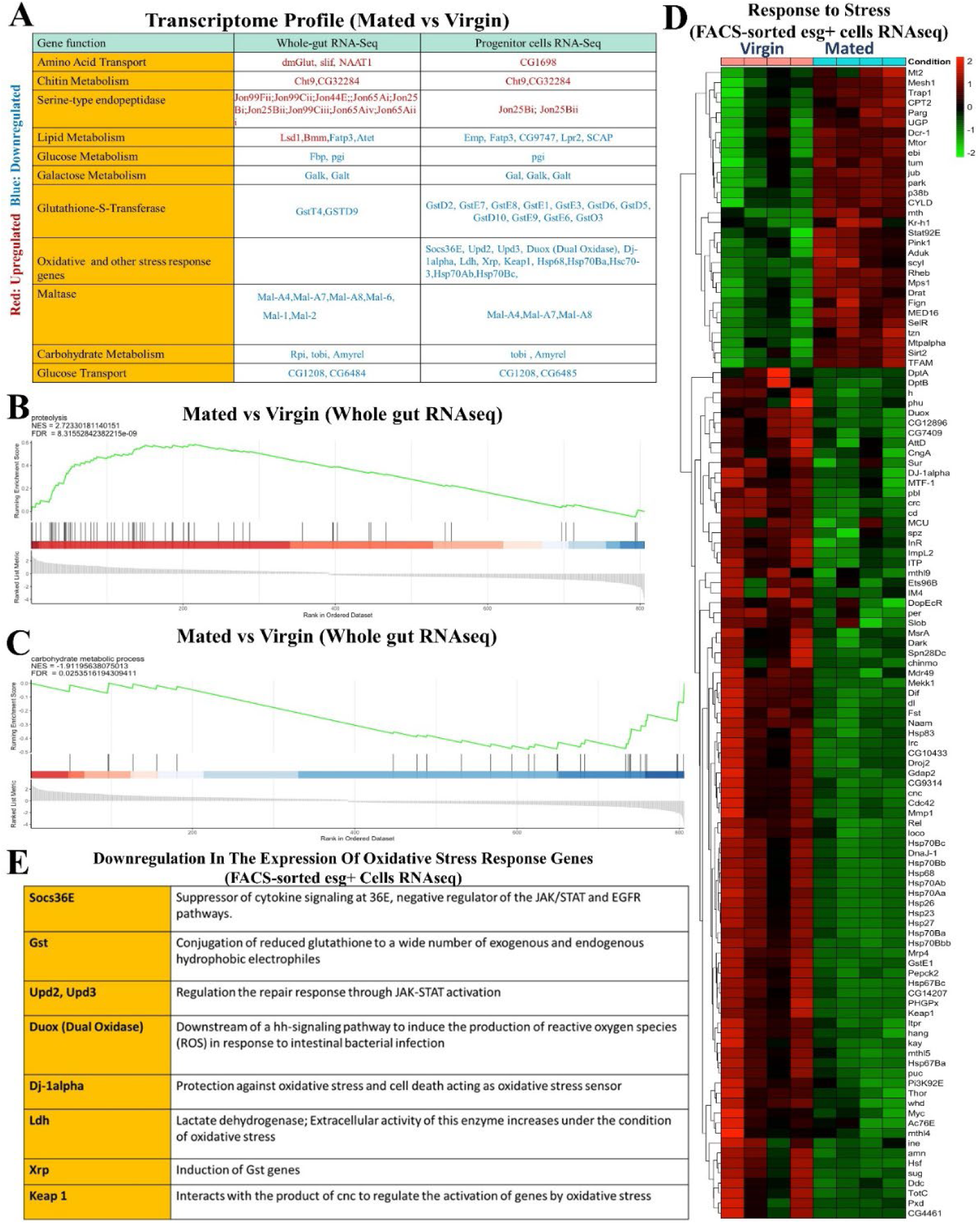
Mated flies have a distinct transcriptomic profile than virgin flies. **A.** Transcript profile comparing gene expression in mated and virgin flies, genes are grouped based on the defined GO terms, genes with transcript upregulation (log-2 fold change ≥1, p-value <0.05) are labeled as red, genes with transcript downregulation (log-2 fold change ≤ -1, p-value <0.05) are labeled as blue. **B-C.** GSEA generated by unbiased hierarchical clustering from whole gut RNAseq shows proteolysis upregulation (B) and carbohydrate metabolic process downregulation (C) in mated flies (*esg-Gal4^ts^*) compared to age and genotype-matched virgin flies. **D.** Targeted heat maps for the genes involved in response to stress (FACS-sorted esg+ cells RNAseq). **E.** List of the genes involved in the oxidative stress response that are downregulated in mated flies *(esg-Gal4^ts^)* compared to age- and genotype-matched virgin flies (FACS-sorted esg+ cells RNAseq).

**Extended Data Figure 2.**
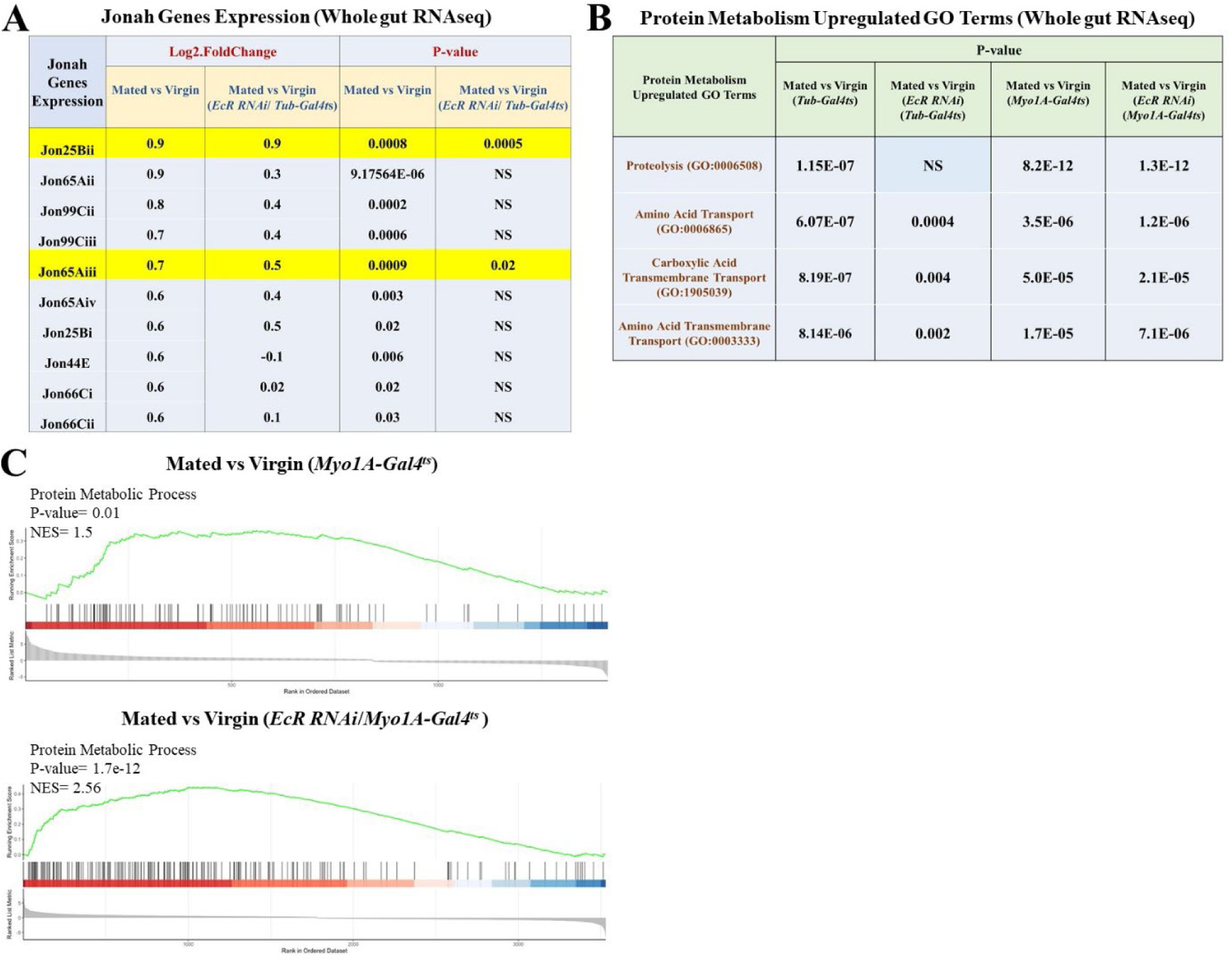
Mating-induced protein metabolism requires EcR. **A.** Jonah gene expression in mated flies compared to virgin controls **+/-** *EcR-RNAi* under the control of the *Tub-Gal4^ts^*, whole gut RNAseq. Each condition had 4 replicates of 15 guts. **B.** Upregulation of protein metabolism GO terms in mated flies compared to virgin controls +/- *EcR-RNAi* under the control of both *Tub-Gal4^ts^* and *Myo1A-* Gal4^ts^, whole gut RNAseq, each/4 replicates/15 guts. **C.** GSEA comparing protein metabolic process genes expression in mated vs. virgin flies +/- *EcR-RNAi* using *Myo1A-Gal4^ts^* driver. The Enrichment Score (ES) measures how much a gene set is overrepresented at either end of a ranking list, while the Normalized Enrichment Score (NES) reflects the degree of this enrichment, shown on the y-axis. GSEA normalizes the ES to account for differences in gene set sizes and their correlations with the expression data.

**Extended Data Figure 3.**
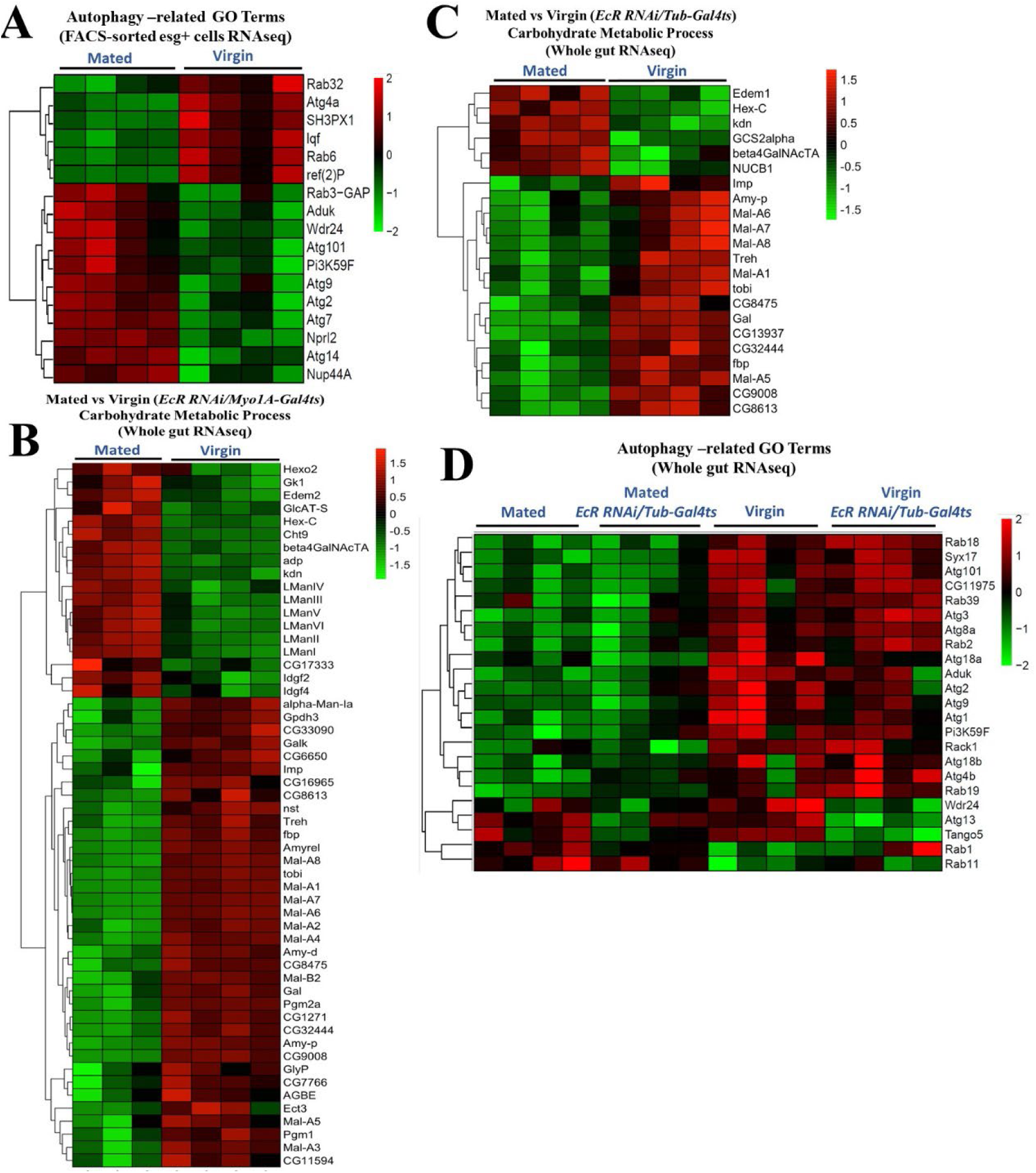
Mating-induced downregulation of carbohydrate metabolism and autophagy gene expression is EcR independent. **A.** Targeted heatmap for the significantly differentially expressed genes involved in autophagy-related GO terms, comparing mated and virgin flies (*esg-Gal4^ts^*) (FACS-sorted esg+ cells RNAseq). **B-D.** Targeted heatmaps for the significantly differentially expressed genes involved in carbohydrate metabolic process (Whole gut RNAseq) comparing mated and virgin flies +/- *EcR-RNAi* using *Myo1A-Gal4^ts^* (B) and *Tub-Gal4^ts^* (C) drivers and comparing autophagy-related genes expression, using *Tub-Gal4^ts^* (Whole gut RNAseq) (D).

**Extended Data Figure 4.**
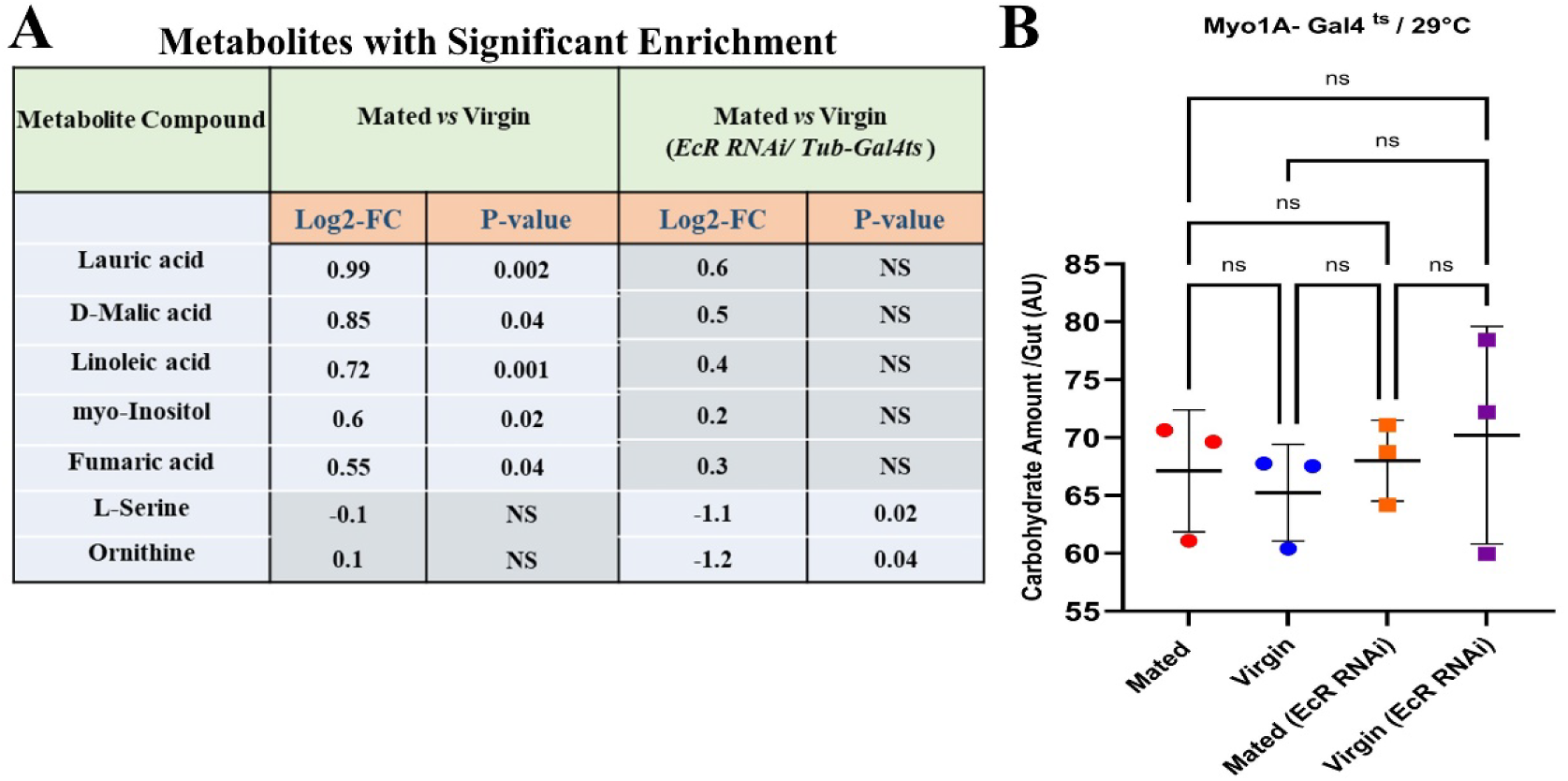
EcR expression modulates the accumulation of fatty acids and TCA cycle metabolites but not carbohydrates in mated flies. **A.** Metabolites with significant enrichment in mated flies versus virgin ones +/- *EcR-RNAi* using *Tub-Gal4^ts^* driver. **B.** Total carbohydrate content in the guts of mated flies and virgin flies +/- *EcR-RNAi* (*Myo1A-Gal4^ts^*) (each/3 replicates/ 15 guts).

**Extended Data Figure 5.**
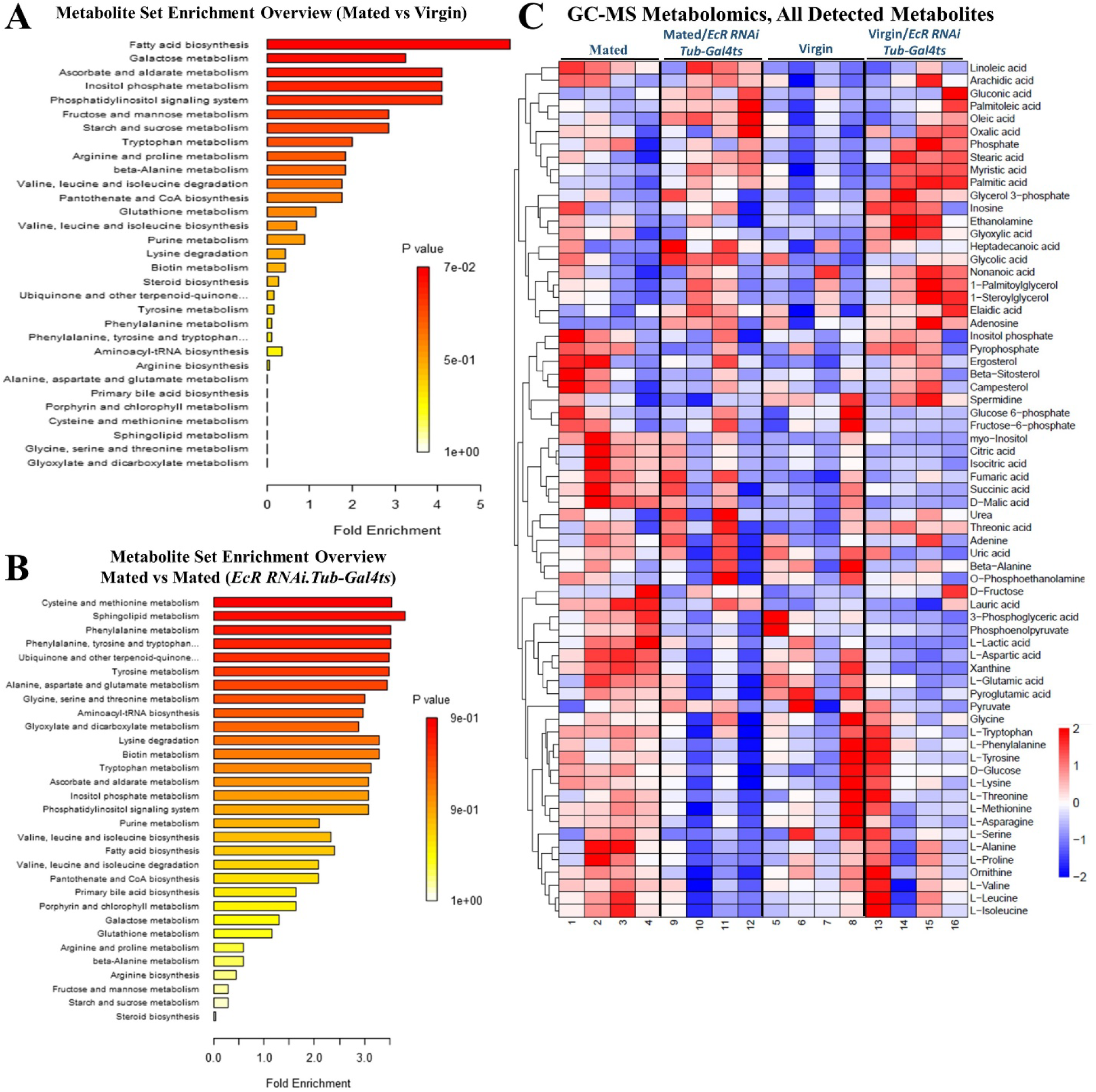
Mating-induced increase in TCA cycle intermediates, and amino acids is associated with EcR expression. Metabolites set enrichment overview, showing the degree of enrichment of different metabolites sets in mated flies (*Tub-Gal4^ts^*) compared to age- and genotype-matched virgin controls **(A)** and compared to mated flies with *EcR-RNAi* expression under the control of *Tub-Gal4^ts^* driver **(B)**; each/ 4 replicates/ 60 guts, the length of each bar shows the degree of enrichment for its correspondent metabolite and the color shows the p-value. **C.** GC-MS data analysis shows all detected metabolite species comparing mated and virgin flies gut +/- *EcR-RNAi* using *Tub-Gal4^ts^* driver depicted in the heat maps based on z-score (each/4 replicates/ 60 guts).

## Notes

### Competing Interest Statement

The authors have declared no competing interest.

